# NR0B2 regulation during Primary Sclerosing Cholangitis defines a metabolic and pre-malignant reprogramming of Cholangiocyte

**DOI:** 10.1101/2021.09.06.459063

**Authors:** Christophe Desterke, Cyrille Feray

**Affiliations:** Centre Hépato-Biliaire, Hôpital Paul-Brousse, Assistance Publique-Hôpitaux de Paris, Université Paris-Saclay, unité Institut National de la Santé et de la Recherche Médicale UA9, Villejuif, France; Centre Hépato-Biliaire, Hôpital Paul-Brousse, Assistance Publique-Hôpitaux de Paris, Université Paris-Saclay, unité Institut National de la Santé et de la Recherche Médicale 1193, Villejuif, France

**Keywords:** cholangiocyte, primary sclerosing cholangitis, metabolism, Farnesoid X receptor, transcriptome, Small Heterodimer Partner, pre-malignancy, TMEM45A

## Abstract

Primary Sclerosing Cholangitis (PSC) is an idiopathic, cholestatic liver disease that is characterized by persistent, progressive, biliary inflammation leading to cirrhosis. These patients present higher risk for developing bile duct cancers.

Biomedical text-mining related to PSC symptoms like: biliary inflammation, biliary fibrosis, biliary cholestasis was initiated to collect gene associations with this pathophysiology. The text mining work was integrated in distinct omics data such as human transcriptome of PSC liver, Farnesoid X receptor (FXR) functional liver transcriptome and liver single cell transcriptome of the Abcb4-/- model of PSC. A molecular network implicated in abnormal hepatobiliary system physiology was built and confirming a major implication of Nr0b2 and its associated nuclear receptors like FXR in a metabolic cascade that could influence immune response. TNFRSF12A/TWEAK receptor, was found up regulated in PSC liver independently of FXR regulation and TWEAK signaling is known for its implication in pre-conditioning niche of cholangiocarcinoma. NR0B2 deregulation in PSC liver was found independent of gender, age and body mass index surrogates. At single cell level, Nr0b2 up regulation was found in cholangiocytes but not in hepatocytes. In affected cholangiocytes, the cell trajectory built on Nr0b2 expression, revealed implication of several metabolic pathways for detoxification like sulfur, glutathione derivative and monocarboxylic acid metabolisms. On this cell trajectory it was discovered some molecules potentially implicated in carcinogenesis like: GSTA3, ID2 and mainly TMEM45A a transmembrane molecule from golgi apparatus considered as oncogene in several cancers. All together, these observations found in humanPSC liver and in its murine models allowed to highlight an independent deregulation of NR0B2 with a metabolic and premalignant reprogramming of cholangiocytes.

## INTRODUCTION

Primary sclerosing cholangitis is a progressive, cholestatic hepatic disease of unknown etiology. It is characterized by progressive inflammation, destruction, and fibrosis of the intrahepatic and extrahepatic bile ducts (Lazaridis & LaRusso, 2016). PSC is a rare disease with an incidence that ranges from 0 to 1.3 cases per 100,000 persons per year (Molodecky et al., 2011). Co-occurrence of inflammatory bowel disease (IBD), most commonly ulcerative colitis (UC), has been reported in up to 80% of patients with PSC (Mertz, Nguyen, Katsanos, & Kwok, 2019). The close association between PSC and IBD indicates the involvement of the gut-liver axis. Also, environmental factor like microbiota has emerged as one of the most important potential environmental players in chronic inflammatory diseases like PSC (Rühlemann et al., 2019). Enterohepatic circulation referred to the metabolite exchange between liver and intestine may play an important aspect of the PSC pathophysiology because metabolome of PSC portal blood and bile samples signed a peculiar molecular profile as compared to ones of disease and non-disease controls (Tietz-Bogert et al., 2018, p.).

PSC causes multifocal strictures of the biliary tree, biliary infections, and accumulation of toxic bile acids, which results in an increased expression of adhesion and antigen-presenting molecules and inflammatory mediators. These observations are a good field for emergence of carcinogenesis that could affect biliary tracts (Chalasani et al., 2000; Rizvi, Eaton, & Gores, 2015). There is no effective treatment which can delay its progression or ameliorate the transplant-free survival. More recently, novel pharmacological agents emerged aiming at: modulation of bile composition, immunomodulation, targeting the gut microbiome, targeting fibrosis (Floreani & De Martin, 2021).

Many aspects of the pathogenesis of PSC remain unclear. Biomedical literature is traditionally used as a way to inform scientists of the relevance of genes in relation to a research topic (Fleuren & Alkema, 2015). Previously, test of distinct biomedical text mining web tools to integrate gene relations with omics data highlighted ‘Génie’ algorithm (Fontaine, Priller, Barbosa-Silva, & Andrade-Navarro, 2011) as a robust and sensitive machine learning tool to rank best literature associated genes. By textmining (Génie) integration in single cell transcriptome it could be possible to highlighted podocytes markers implicated in Focal Segmental Glomeral Sclerosis (Desterke, Lorenzo, & Candelier, 2021). ‘Génie’ algorithm also contributed to understand lipidome deregulations during nonalcoholic fatty liver disease (Pirola & Sookoian, 2021). ‘Génie’ is a powerful and fast tool that prioritizes the whole gene sets of hundreds of species for any biomedical topic, taking advantage of annotations-linking gene and bibliography (Fontaine et al., 2011). In this work, biomedical text-mining related to PSC symptoms allowed to enrich a gene set of molecules that was used originally to analyze different omics data coming from human PSC liver biopsies and distinct animal models of PSC. Independent deregulation of NR0B2 in PSC allowed to highlight a metabolic and pre- malignant reprogramming of the affected cholangiocytes.

## MATERIAL AND METHODS

### Biomedical Pubmed textmining

‘Génie’ Data mining tools performed gene prioritization (Fontaine et al., 2011) by linking scientific litterature from MEDLINE database with gene information from NCBI Gene database and orthologue gene information from different species with HomoloGene database (E. Sayers, 2010; E. W. Sayers et al., 2009). Génie algorithm works by connecting gene identifier and gene orthologue NCBI databases with genes related to input keywords obtained by building a textmining classifier with article abstracts from PUBMED literature ressource. Primary Sclerosing Cholangitis related symptom Medical Subject Headings (MeSH) terms were individually submitted to ‘Génie’ algorithm on 2021-july-2nd. All abstract of PUBMED were taking as background for the analysis of genes corresponding to Homo sapiens coding proteins. All orthologue information were collected during the process, p-value cutoff for abstract selection was fixed to p<0.01 and False Discovery Rate cutoff for gene selection was fixed to p<0.01. This last step of FDR adjustment allowed to correct false positive discovery rate and so to minize the influence of one pecular article in the globality of the analysis of gene enrichment.

### Public datasets

#### Transcriptome studies

1/ Human liver biopsies (n=134), GSE61256 (Horvath et al., 2014): transcriptome experiments performed on human biopsies were downloaded and annotated with the corresponding annotation platform: GPL11532 used for technology version DNAChipAffymetrix Human Gene 1.1 ST Array (alias HuGene-1_1-st) available at the fellowing address: https://www.ncbi.nlm.nih.gov/geo/query/acc.cgi?acc=GPL11532 (accessed on 2021, july 2^nd^).

2/ Mus musculus liver FXR functional study GSE54557 (Zhan et al., 2014): C57BL/6J Mus musculus liver samples of 10-16 weeks. Transcriptome of liver samples in presence or not of liver FXR knock-out and with or without treatment with GW4064 FXR agonist experience en triplicate. Transcriptome matrix was downloaded and annotated with the annotated platform GPL8321 corresponding to the DNAchip technology: Affymetrix Mouse Genome 430A 2.0 Array (alias Mouse430A_2) available at the following address: https://www.ncbi.nlm.nih.gov/geo/query/acc.cgi?acc=GPL8321 (accessed on 2021, july 2nd).

### Single cell transcriptome study

Mus musculus Abcb4-/- single cell transcriptome liver samples GSE168758 (Reich et al., 2021): Single cell transcriptome libraries of liver cells were prepared with Chromium Single Cell 3’ NextGEM Reagent Kit v3.1 (10X Genomics) to performed sequencing on NextSeq 550 (Illumina) and demultiplexing pipeline with CellRanger v5.0.0 (10x Genomics). MTX matrix file format were downloaded at the following address: https://www.ncbi.nlm.nih.gov/geo/query/acc.cgi?acc=GSM5165876 (accessed on 2021, july 2nd).

### Bioinformatics analyses

Bioinformatics analysis were performed in R software environment version 4.1.0 under Ubuntu version 20.04 LTS. Graphical representation were performed with ggplot2 graph definition implemented in ggplot2 R-package version 3.3.5 (Wickham, 2009, p. 2).

### Transcriptome Analyses

Unsupervized analysis performed by principal component analysis used FactoMineR R-package version 2.4 (Lê, Josse, & Husson, 2008). Gene Functional enrichment analyses were performed with Toppgene web application (J. Chen, Bardes, Aronow, & Jegga, 2009). Differential expressed genes were identified with LInear Model from Microarray (LIMMA) algorithm implemented in R Bioconductor package version 3.44.3 (Ritchie et al., 2015). FXR regulation dependency analysis with Pavlidis Template Matching algorithm with retaining significant genes on absolute value of correlation coefficient and with a cutoff p-value less than 0.05 (Pavlidis & Noble, 2001). Expression heatmap were drawn with pheatmap R-package version 1.012.

### Single cell transcriptome analysis

Single transcriptome analyses at whole liver level were performed with Seurat4 R-package version 4.0.3 (Butler, Hoffman, Smibert, Papalexi, & Satija, 2018). Briefly, after canonical correlation, filtration (cells expressing minimally 300 genes) and scaling between replicated samples (n=4) of wildtype and Abcb4-/- groups a normalized single cell object comprising 46.087 transcriptomes was built. Dimension reduction was carried out by principal component analysis on most variable features (50 components) and secondly: t-distributed stochastic neighbor embedding (t-SNE) and Uniform Manifold Approximation and Projection (UMAP) dimensionality reduction was performed on the 30 best components of the PCA. Cell cluster communities were detected graph-based clustering approaches (Xu & Su, 2015). Abcb-/- cholangiocytes identified in clusters 0-5-11 were select were randomly down sampled to 1/10e of their initial amount to build a matrix finally containing 1316 cells. Single cell trajectory performed on Abcb4-/- cholangiocyte with monocle2 R Bioconductor package version 2.20.0 (Trapnell et al., 2014). Cell hierarchy was construct on alternative expression of Sox9 and Nr0b2 in Abcb4-/- cholangiocytes. Pseudotime transformation on cell trajectory was performed with DDRtree algorithm and pseudotime cluster identification was done by pseudotime expression heatmap drawn with best genes found significant

### Network analyses

Gene and function relations identified were collected with functional enrichment and network were built with Cytoscape standalone software version 3.6.0 (Cline et al., 2007) for their visualization. Protein-protein interaction (PPI) network were performed with NetworkAnalyst (Xia, Gill, & Hancock, 2015) webtools based on interaction network databases: STRING (Szklarczyk et al., 2017) or liver specific DifferentialNet. Functional inference was done on PPI networks with Gene Ontology Biological Process (GO-BP) database (The Gene Ontology Consortium, 2017).

## RESULTS

### Biomedical text mining for discovery of gene relations with symptoms of primary sclerosing cholangitis

Classically, the litterature dealing with genes, as stored in the MEDLINE database (E. W. Sayers et al., 2009) of biomedical references has been used to do gene prioritization. Many methods for gene prioritization using litterature data are based on co-occurence analysis of keywords and extract gene names from abstracts (Krallinger, Valencia, & Hirschman, 2008). ‘Génie’ algorithm is based on Naive bayes classifier and take advantage of linking orthologues genes study in many different species (Fontaine et al., 2011). Primary Sclerosing Cholangitis (PSC) is an Idiopathic, Heterogeneous, cholestatic liver disease that is characterized by persistent, progressive, biliary inflammation and fibrosis (Lazaridis & LaRusso, 2016). Three related MESH terms for indirect characterization of PSC symptomatology were used to process in ‘Génie’ algorithm and trigger gene related to the disease with the following exact terms: ‘biliary inflammation’, ‘biliary fibrosis’, ‘biliary stasis’. These respective analyses allowed to collect a common set of 525 ranked genes (workflow, **Figure 1** and **Supplemental Table 1**). NR1H4 alias FXR Farnesoid X receptor was found as the best ranked gene at cross of these three independent text mining analyses (**Supplemental Table 1**). This text mining results allowed to give us a restrictive gene list (525) to investigated downstream omics analyses in context of PSC disorder as it is highlight graphically on the workflow of the article (**Figure 1**). Starting omics analyses with reduced size gene sets allowed to drastically eliminate background noise and False Discovery Rate in these experiments as it is shown with Gene Set Enrichment Analysis (GSEA) procedure (Subramanian et al., 2005).

**Figure 1:**
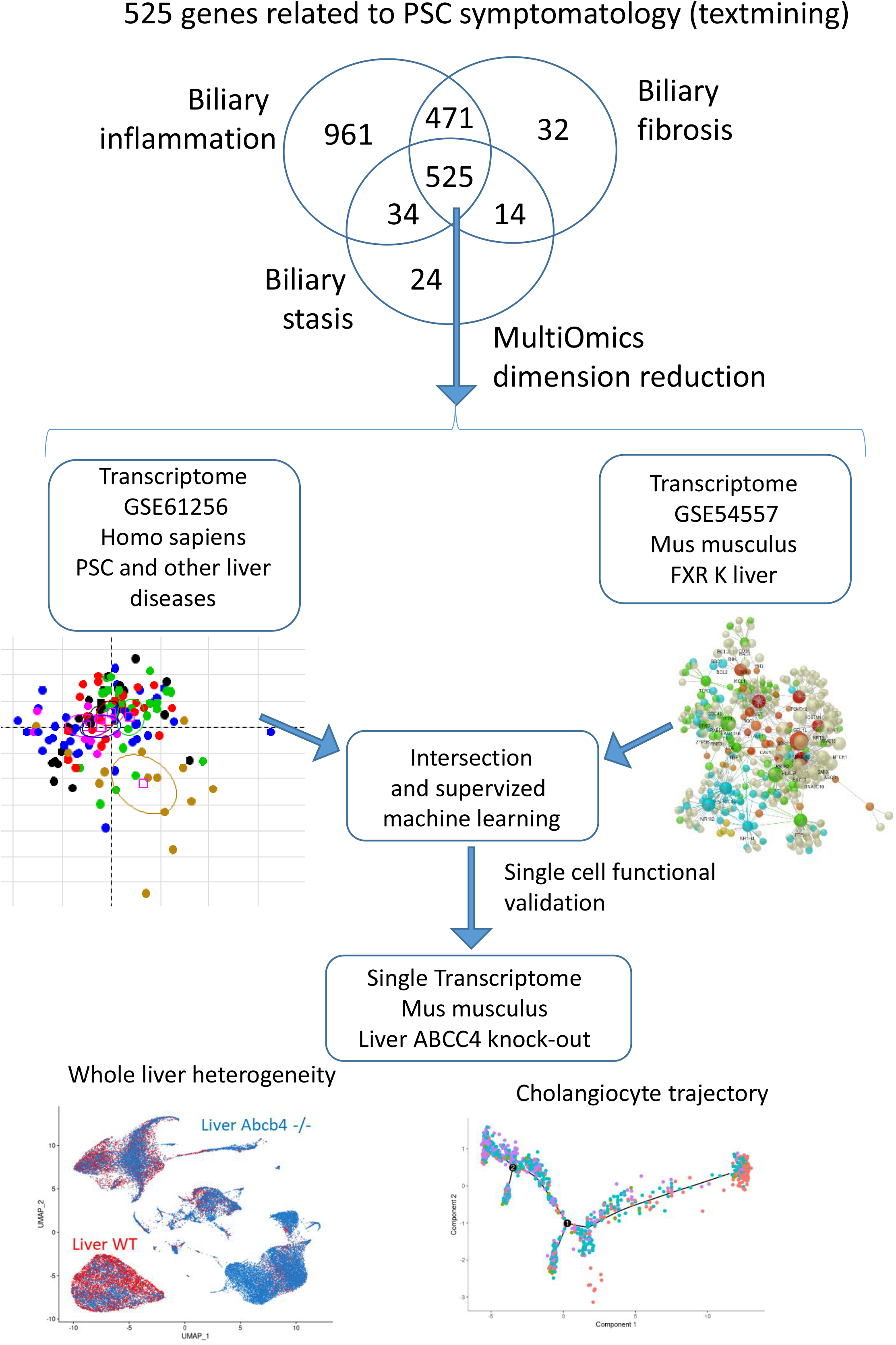
workflow of bioinformatics analyses: this figure describes the workflow of the bioinformatics process performed on data used in this work.

### Abnormalities in physiology of hepatobiliary system during primary sclerosing cholangitis

Primary sclerosing cholangitis (PSC) is a chronic hepatic syndrome with fibro-inflammatory affection of the bile ducts. In order to test our txt mining PSC symptom related gene set, a bioinformatics investigation was performed on transcriptome study GSE61256 (**Figure 1**) (Horvath et al., 2014) which is composed of liver biopsy samples from PSC (n=14) patients as compared to distinct control groups such as: PBC (n=11), normal control (n=38), normal obese (n=24), NAFLD (n=23), NASH (n=24). Data from GSE61256 were reduced in dimension with the 525 PSC symptom related genes but only 507 of them were find provided in these type of experiments. Unsupervised principal component analysis revealed an individual stratification of PSC samples as compared to every other control groups and especially as compared to PBC control which could harbored also these three phenotypes (p-value=6.89E-4, **Figure 2A**). With less importance, these genes also discriminates PBC samples from other liver control groups in the second map of PCA analysis (Supplemental Figure 1). These results suggested that in its entirety PSC symptom related gene set obtained by text mining seems to be specific to predict liver sample transcriptome of PSC as compared to divers control groups. Effectively, starting from these 507 molecules supervised ‘LInear Model from Microarray’ (LIMMA) algorithm for differential expression analysis allowed to characterized a significant expression profile of PSC liver samples as compared to all other pooled control groups (**Figure 2B** and **Supplemental Table 2**). Among the best up regulated genes, FOXP3 was found the best one followed by NR0B2, TNFRSF1A and FGF19. A distinct trait of expression of up regulated for PSC samples could be observed on the right of expression heatmap performed with up regulated genes (**Figure 2C**). Functional enrichment performed with these PSC up regulated genes on Mouse Phenotype database allowed to characterized were found mainly implicated in abnormal liver physiology and abnormal hepatobiliary system physiology (**Figure 2D**). As PSC is a hepatic disorder affecting bile ducts a functional enrichment network was built on affection of hepatobiliary system. On this hepatobiliary system phenotype network we could observed that some of the top up regulated genes in PSC are present such as: FOXP3, TNFRSF12A, TNFRSF1A, LCN2 and NR0B2 (**Figure 2E**). Starting from PSC symptom related gene found by text mining we could predict with liver samples, a PSC specific up regulated network implicated in hepatobiliary system phenotype.

**Figure 2:**
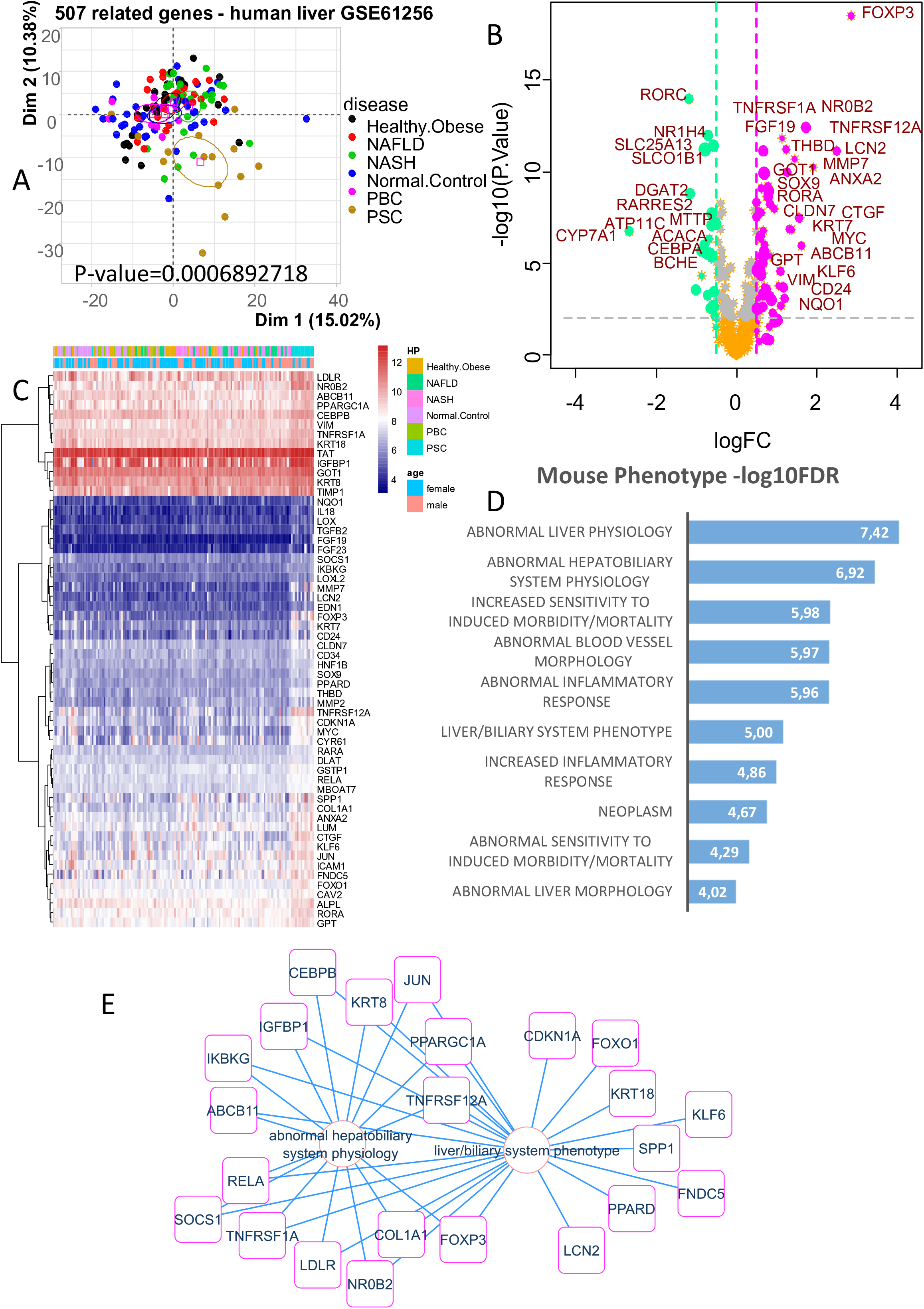
Abnormal hepatobiliary system physiology in liver biopsies of Primary sclerosing cholangitis liver biopsy: from A to E: investigations on transcriptome dataset GSE61256; A/ principal component analysis performed with 507 detected symptom related genes on liver biopsy transcriptome from distinct group of diseases and controls, PSC: primary sclerosing cholangitis, PBC: primary biliary cholangitis, NAFLD: Nonalcoholic fatty liver disease, NASH: Nonalcoholic steatohepatitis; B/ Volcanoplot of differential expressed genes (DEGs) between PSC liver and other liver biopsies, C/ Expression heatmap of significant up regulated genes in PSC liver samples; D/ Functional enrichment performed on Mouse Phenotype database with genes up regulated in PSC liver samples; E/ Functional enrichment network with PSC up regulated genes and implicated in hepatobiliary mouse phenotype.

### Hepatic regulation of immune response and intracellular receptor signaling under FXR dependency

In PSC symptom related genes found by text-mining we could observed that NR1H4 alias Farnesyl X receptor (FXR) was the best ranked gene associated to the literature in relations with input symptom MeSH. FXR is a nuclear receptor implicated in bile acide metabolism and have been widely study in hepatocyte biology such as whole gene ChIP-sequencing experiments (Zhan et al., 2014). Liver transcriptome study were also performed on hepatic functional studies targeting FXR: such as GSE54557 dataset which investigate liver samples of Mus musculus with or without liver FXR knock-out reversed of not by GW4064 agonist treatment (Zhan et al., 2014). Data from GSE61256 were reduced in dimension with the 525 PSC symptom related genes and Pavlidis Template Matching (PTM) algorithm was used to understand FXR regulation dependency: FXR liver knock-out reversed by GW4064 FXR agonist (**Figure 3A** and **Supplemental Table 3**): 91 genes were found significantly regulated through FXR dependency in liver samples. These significant genes allowed to significantly stratified experimental samples of FXR liver KO from wildtype ones (WT) and reversion by FXR agonist was obtained only in wildtype samples (p-value=1.66.E-4, **Figure 3B**). Molecules with positive regulation following FXR liver dependency were selected to draw an expression heatmap (Figure 3C) which implicated a protein-protein interaction (PPI) network mainly implicated in regulation of immune response (nodes in green, **Figure 3D**) and in intracellular receptor mediated signaling pathway (in blue, **Figure 3D**). Concerning intracellular receptor signaling, three main blue crosstalk could be observed on the PPI network (**Figure 3D**): NR1H4, NR1I2 and PPARGC1A which are closely connected. Concerning immune response we could observed (**Figure 3C**) a liver FXR regulation dependency of CD274 alias PDL1 immune checkpoint. These results suggest that starting from PSC symptom related gene list we could predict a hepatic expression profile under FXR regulation dependency and so confirming its potential functional importance in the pathology.

**Figure 3:**
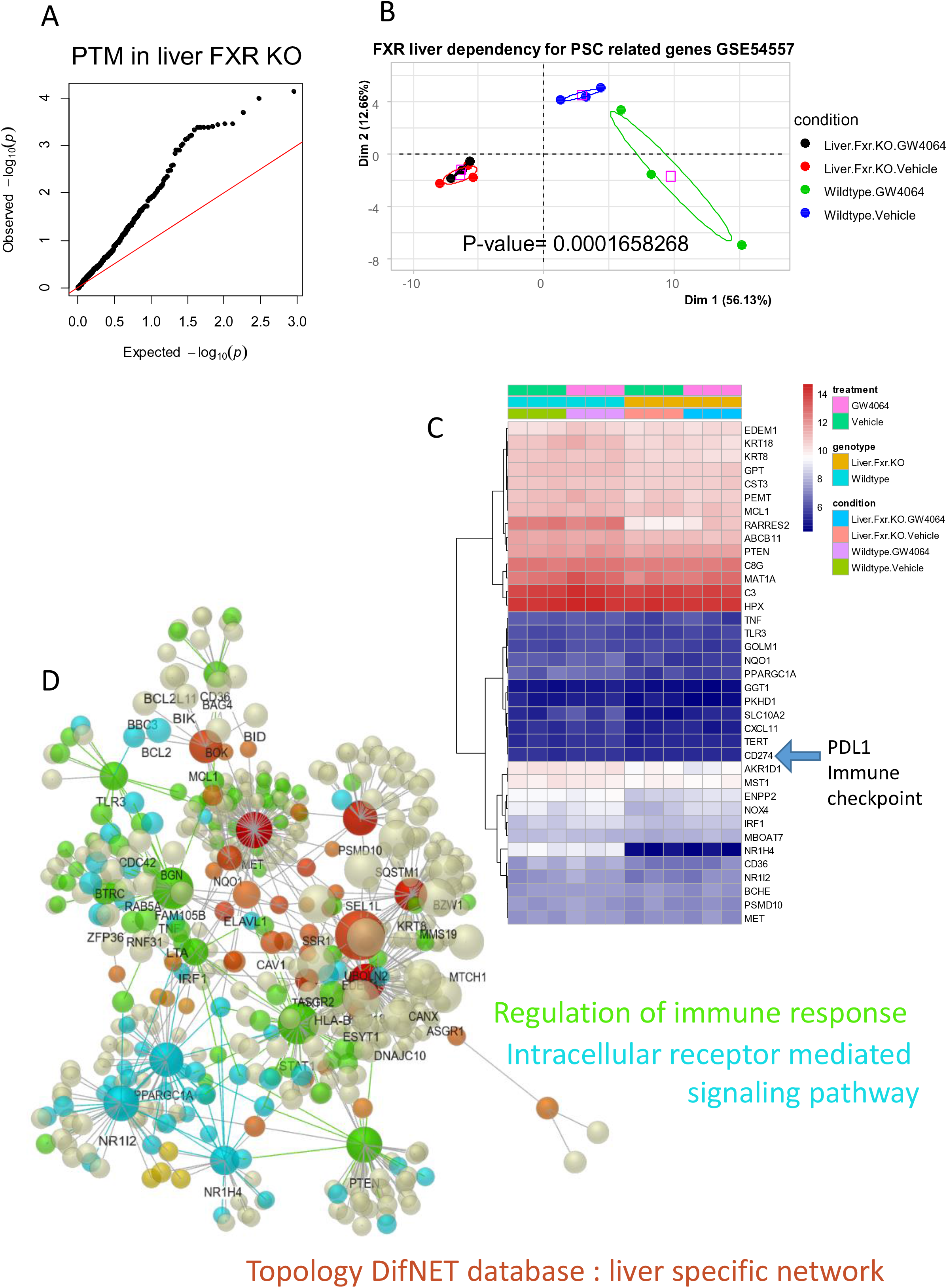
Hepatic regulation of immune response and intracellular receptor signaling under FXR dependency: from A to D: investigations on transcriptome dataset GSE54557; A/ qqplot of p-values obtained by Pavlidis Template Matching (PTM) algorithm with symptom related genes for liver FXR regulation dependency; B/ Principal component analysis performed with PSC symptom related genes under liver FXR regulation dependency, KO: knock-out, GW4064: FXR agonist; C/ Expression heatmap for PSC symptom related genes found positively correlated to hepatic FXR regulation; D/ Protein-protein interaction network based on PSC symptom related genes up regulated in PSC liver samples with colored function inference of Gene Ontology Biological Process (GO-BP) database: in green enrichment of immune response function and in blue intracellular receptor signaling function.

### Nrb02 is the best predictive marker regulated in liver of primary sclerosing cholangitis under FXR dependency

In order to evaluate FXR dependency in PSC related human liver expression profile that we identified we investigated to cross the human study signature (**Supplemental Table 2** and **Figure 2**) with liver FXR functional signature (**Supplemental Table 3** and **Figure 3**). This investigation allowed to highlight 34 common regulated molecules between the two studies (**Figure 4A**). Proceeding supervised machine learning by leave one out cross validation algorithm (pamr) between human liver PSC samples and other liver control samples (GSE61256), it could possible to defined a predictive molecular ranking (**Supplemental Table 2**) with a minimal misclassification error of 8.9 percent between samples (**Figure 4B**). Inside these best molecular ranking to predict human PSC liver we could observed some molecules with an FXR dependent regulation (in pink, **Figure 4C**), such as: NR0B2, CYP7A1, DGAT2, NR1H4 (alias FXR), TNFRSF1A, BCHE, KLF6 and CEBPA. NR0B2 at the top of PSC prediction with FXR dependency was investigated by univariate analysis stratified on gender of the patients. Effectively, NR0B2 was found up regulated in PSC samples and more sensitively in samples of female gender (**Figure 4D**). In a biological and clinical multivariate model, we could observed that up regulation of NR0B2 was significantly up regulated in PSC liver samples but independently of the gender, the age and body mass index (BMI) of the patients. These results suggest that the nuclear receptor NR0B2 is the best predictive marker up regulated in PSC liver which harbored an FXR dependent regulation is its regulation is independent of the age, gender and BMI of the patients.

**Figure 4:**
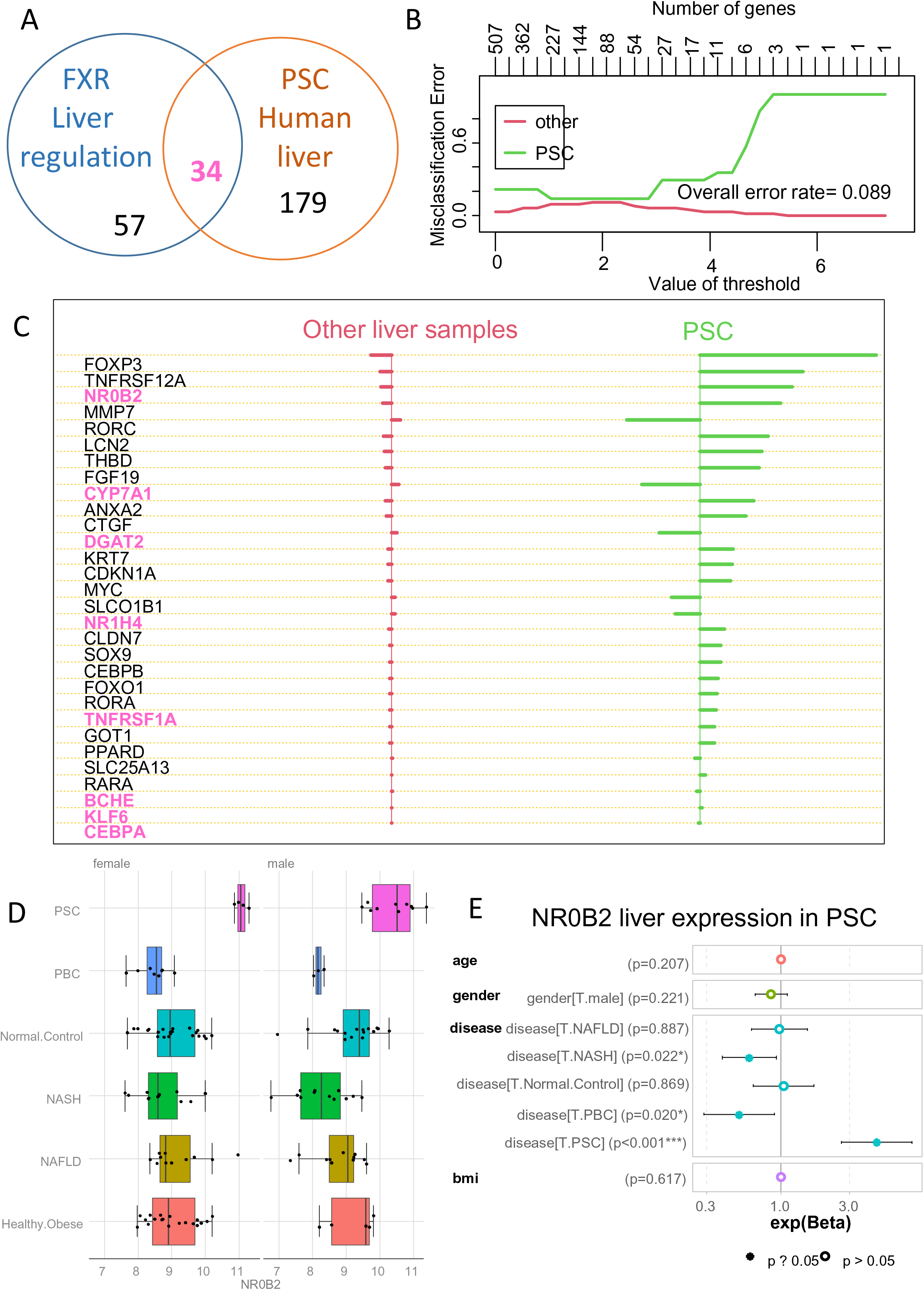
Nrb02 is the best predictive marker regulated in liver of primary sclerosing cholangitis under FXR dependency: A/Venn diagram merging FXR liver regulation dependency of PSC related genes found up regulated in PSC human liver biopsies; B/ Supervised machine learning misclassification plot by class obtained with PSC symptom related genes to stratified PSC liver samples from other control samples in GSE61256; C/ Ranked predictive scores obtained by machine learning on GSE61256 dataset with PSC symptom related genes, FXR hepatic regulation dependency is highlighted in pink for concerned genes; D/ Boxplot of Nr0b2 expression regulation through liver group samples and stratified on gender (GSE61256); E/ Multivariate model based on Nr0b2 expression: exponential coefficients and their confident intervals (95 percent) were drawn for each covariates.

### Up regulation of Nrb02 in cholangiocyte compartment of Abcb4-/- liver

Abcb4-/- (Mdr2) mice are widely used as model for sclerosing cholangitis because it recapitulates the progressive fibrosing cholangitis aspect of human PSC and also develop intrahepatic malignancy (Fickert et al., 2004, 2014). Single cell transcriptome performed in liver of wild type and Abcb4-/- mice models have been provided in GSE168758 dataset (Reich et al., 2021, p. 5). With 4 replicates of liver in groups WT and Abcb4-/- a single cell normalized object was built with 46.087 cells which individually expressed a minimum of 300 genes. After dimension reduction and clustering 15 cell communities were identified and classed in five main cell types: hepatocytes, cholangiocytes, endothelial cells, macrophages and lymphocytes (**Figure 5A** and **Supplemental Figure 2**). Wild type livers harbored more hepatocytes as compared Abcb4-/- livers (in red, **Figure 5B)** and Abcb4 livers harbored more cholangiocytes than wild type livers (in blue, **Figure 5B**). Sox9 is a transcription factor implicated in the biliary development (Antoniou et al., 2009). Epcam and Sox9 expressions follow CK19 expression during human intrahepatic biliary tree formation through several developmental stages involving an initial transition of primitive hepatocytes into cholangiocytes (Vestentoft et al., 2011). Sox9 was found over expressed in Abcb4-/- livers as compared to wild type ones (**Figure 5C**) and especially in cholangiocyte compartment (**Figure 5D**) which correspond to cell clusters : 0, 5 and 11 (**Supplemental Figure 3A**). Similarly, an overexpression of Nr0b2 was found in Abcb4-/- livers as compared to wild type ones (Figure 5E). Expression of Nr0b2 was found detected in hepatocyte and cholangiocyte cell compartment in both liver genotypes (**Figure 5F**). Overexpression of Nr0b2 in liver Abcb4-/- livers concerned cell clusters 0, 5 and 11 (**Figure 5G and Supplemental Figure 3B**) which were confirmed as expressing Sox9 and Epcam cholangicoytes markers (**Supplemental Figure 3A and 3C**). Absence of Nr0b2 regulation was observed in hepatocyte compartment of Abcb4-/- (cluster 1, **Figure 5G**). These results suggest that Nr0b2 nuclear receptor was found up regulated in cholangiocyte compartment of the Abcb4-/- liver: mice model of sclerosing cholangitis.

**Figure 5:**
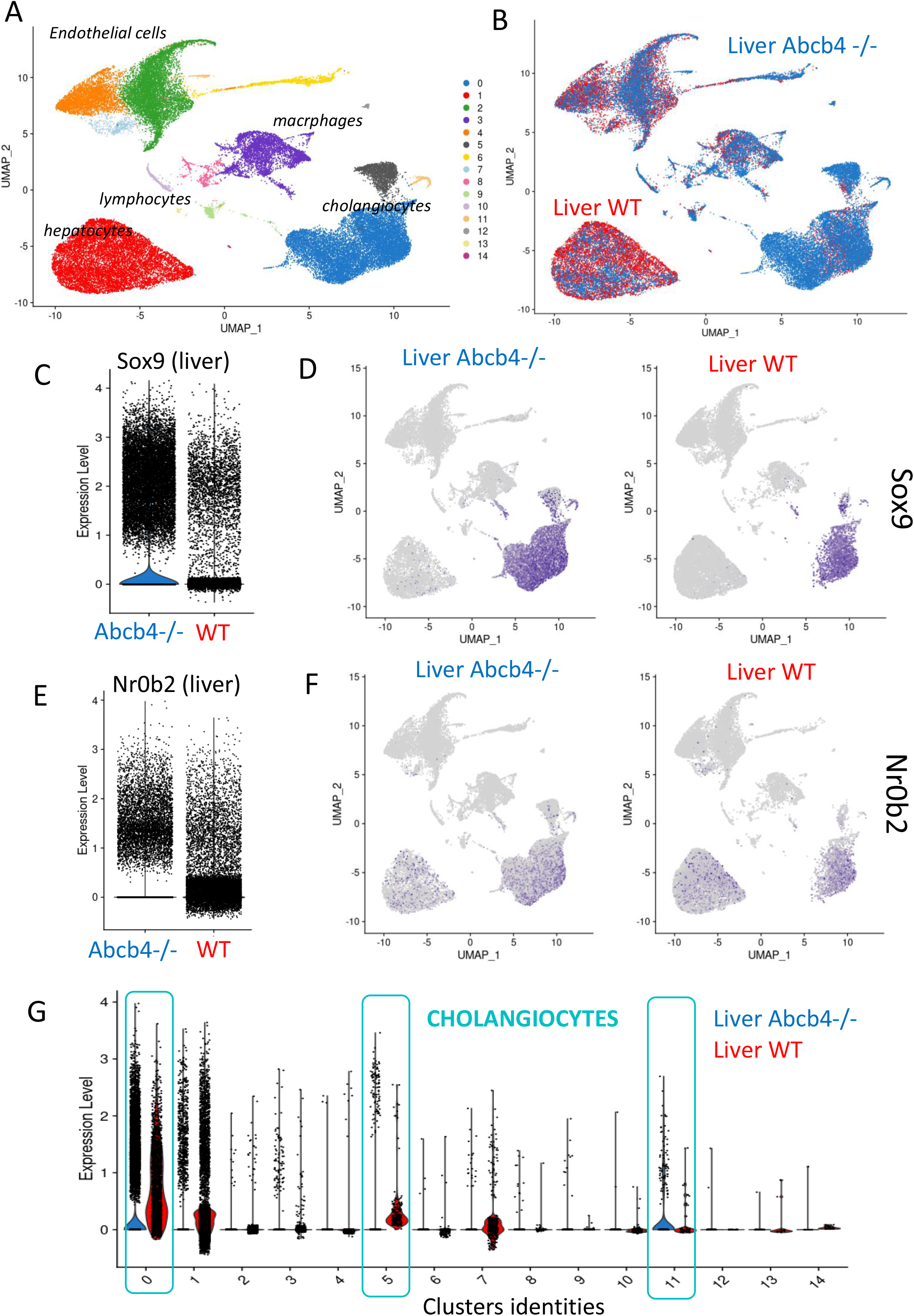
Up regulation of Nrb02 in cholangiocyte compartment of Abcb4-/- liver: from A to G: investigations on whole liver single cell transcriptome dataset GSE168758; A/ Uniform Manifold Approximation and Projection (UMAP) dimension reduction with identification of the distinct cell compartments in whole liver samples: colors reflect 15 distinct cluster cell communities; B/ UMAP with genotype stratification: WT: wildtype, Abcb4-/-: Abcb4 knock-out; C/ Violinplot of Sox9 expression in whole liver stratified on genotypes; D/ Sox9 feature plot stratified on genotypes: blue for positive expression; E/ Violinplot of Nr0b2 expression in whole liver stratified on genotypes; F/ Nr0b2 feature plot stratified on genotypes: blue for positive expression; G/ violin of Nr0b2 expression stratified on cluster cell communities and genotypes.

### Metabolic regulation during cell trajectory of Abcb4-/- cholangiocytes

Nr0b2 alias small heterodimer partner is a nuclear receptor identified as a key transcription regulator affecting divers functions implicated in metabolism such as: bile acid metabolism, lipid metabolism, glucose and energy homeostasis (Zhang, Hagedorn, & Wang, 2011, p. 2010). Nr0b2 was found as best predictive marker of PSC in liver with FXR regulation dependency but its overexpression in Abcb4-/- mice model affect cholangiocyte but not hepatocyte cell compartment. Starting from single cell transcriptome of GSE168758 dataset, a random down sampling subset of Abcb4-/- cholangiocytes was selected to build a single cell trajectory based on alternative expression of Nr0b2 and Sox9. With an affected cholangiocyte compartment comprising in total 1316 cells, 238 cells were characterized as Sox9 Nr0b2 double negative, 51 cells were characterized as negative for Sox9 and positive for Nr0b2, 406 cells were characterized as positive for Sox9 and negative for Nr0b2, and 621 cells were characterized as Sox9 and Nr0b2 double positive (**Figure 6A**). The alternative expression of Sox9 and Nr0b2 reflect a cell heterogeneity in Abcb4-/- the cluster heterogeneity of cholangiocytes clusters 0, 5 and found previously found at whole liver level (**Supplemental Figure 4A and 4B**). Pseudotime transformation of single cell transcriptome in this affected compartment (**Figure 6** and **Supplemental Figure 4C**) reflect a cell trajectory with progressive modification of Sox9 (**Supplemental Figure 4D**) and Nr0b2 (**Supplemental Figure 4E**) expressions. Identification of most significant regulated genes on this cell trajectory (**Supplemental Table 4**) allowed to identified three distinct regulated gene clusters (**Figure 6C**). Expressions of Sox9 and Nr0b2 were found closely regulated on the trajectory of the pseudotime green gene cluster (**Figure 6C**). Among the pseudotime green cluster, expression of Alb, Gsta3, Id2 and Tmem45a were found closely regulated with Nr0b2 and Sox9, especially Gsta3 very near Nr0b2 pseudotime regulation (**Figure 6C** and **Supplemental Figure 4F**). Protein-protein network performed with genes belonging to green pseudotime gene cluster with functional inference of Gene Ontology Biological Process database allowed to identified a main functional enrichment of positive regulation of metabolism (False Discovery Rate p-value=5.2E-23, Figure 6D). This metabolic functional enrichment allowed to mainly implicated sulfur compound metabolism, glutathione derivative biosynthesis, cellular detoxification and monocarboxylic acid metabolism (**Figure 6E**) which share some main regulated molecules (**Figure 6F**). These results suggest that heterogeneity of Nr0b2 expression inside Abcb4-/- cholangiocyte affected compartment defined a cell trajectory implicating an important regulation of metabolism with detoxification process.

**Figure 6:**
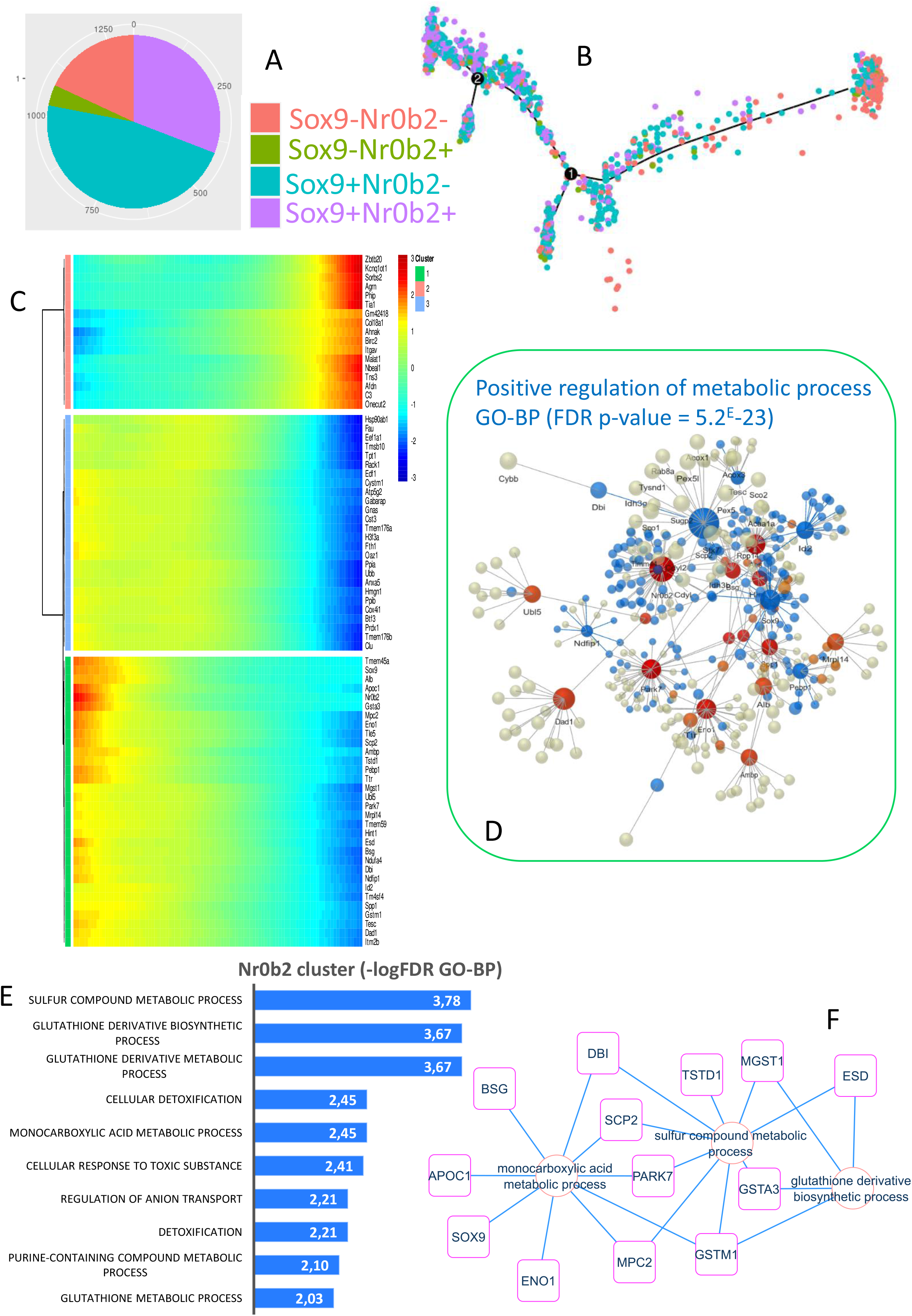
metabolic regulation during cell trajectory of Abcb4-/- cholangiocytes: from A to F: downsampling of Abcb4-/- identified cholangiocytes in liver single cell transcriptome dataset GSE168758; A/ Pie chart of cell stratification depending of alternative expression of Nr0b2 and Sox9 in cholangiocyte of Abcb4-/- liver samples; B/ Abcb4-/- cholangiocyte single cell trajectory identified on altenative expression of Nr0b2 and Sox9; C/ Pseudotime expression heatmap of genes found significantly regulated on Nr0b2-Sox9 single cell trajectory of Abcb4-/- cholangiocytes: green cluster follow Sox9 and Nr0b2 pseudotime expression; D/ protein-protein interaction network identified on pseudotime green cluster: blue color characterized network inference of positive regulation of metabolism process; E/ Barplot of functional enrichment performed on pseudotime green cluster with Gene Ontology Biological Process database, FDR: False Discovery Rate; F/Functional enrichment network performed with best metabolism function enriched on pseudotime green cluster.

## DISCUSSION

Primary sclerosing cholangitis is a chronic, fibro-inflammatory disorder affecting intra/extrahepatic bile ducts without precise etiology and efficient therapy. The resulting cholestasis may progess to cirrhosis and liver failure (Karlsen, Folseraas, Thorburn, & Vesterhus, 2017). Enterohepatic circulation have been proposed to participate to the pathophysiology of this disease because it could be diagnose a preponderance of inflammatory bowel disease in PSC patients (Boonstra et al., 2012) and also it have been discovery that the gut microbiota of PSC patients is distinct from those inflammatory bowel disease patients and healthy controls (Sabino et al., 2016). Progress remains to be made in the understanding of the physiopathology of this disease in order to elucidate why some of these patients have a higher risk of developing cholangiocarcinomas (Rizvi et al., 2015). Few liver transcriptome experiments were available in PSC disease (Horvath et al., 2014): experiments in which it might be interesting to observe cellular mechanisms consecutives to the symptoms of PSC disorder. The integration of biomedical text mining into high-dimensional omics data such as transcriptome data drastically reduces the error of positive false discovery (Desterke et al., 2021). With the aim of increasing the authenticity of discovery of these mechanisms in human liver transcriptome samples from PSC, a biomedical text-mining pipeline has been developed based on a triad of symptoms characterizing PSC: biliary inflammation, biliary fibrosis and biliary stasis (Lazaridis & LaRusso, 2016). The intersection of these text-mining researches have made it possible to link 525 genes to the symptomatic of PSC disorder. The best literature associated gene by text-mining was NR1H4 alias FXR Farnesoid X receptor a nuclear receptor implicated in bile acid metabolism. Nuclear receptors (NRs) are a unique family of transcription factors (TFs) that are being recognized as key regulators of multiple functions in almost all aspects of mammalian development, metabolism and physiology (Gronemeyer, Gustafsson, & Laudet, 2004). FXR is highly expressed in liver and is used as bile acid sensor by regulating genes that are critically involved in bile acid homeostasis, including bile acid biosynthesis, conjugation, and entero-hepatic circulation (Sinal et al., 2000). PSC symptom related genes found by biomedical text mining were used to analyzed human liver transcriptome and predicted a stratification of PSC samples as compared to samples from different controls: healthy normal liver, obese healthy donors, NAFLD, NASH and PBC. A PSC expression profile was able to be identified by differential expression analysis which highlighted important over expression of distinct molecules. FOXP3 was found as the best overexpressed gene in liver of PSC patients. FOXP3 transcription factor characterized the phenotype of regulatory T-lymphocyte immune cell sub population (Sakaguchi, Miyara, Costantino, & Hafler, 2010) and bile acid metabolites have been shown to be implicated in the control regulatory T-lymphocyte cell differentiation (Hang et al., 2019). In genome of PSC patients, polymorphisms found in IL2RA gene affected FOXP3+ regulatory T-cell subpopulation of these patients (Sebode et al., 2014). Among best up regulated genes found in PSC it was found NR0B2 alias small heterodimer partner (SHP). NR0B2 has been identified as key transcription regulator implicated in divers biological function such as: bile acids; cholesterol and lipid metabolism, glucose and energy homeostasis (Zhang et al., 2011). NR0B2 alias Small heterodimer partner (SHP) is a unique nuclear receptor (NR) distinct from other conventional NRs in both its structure and function. The presence of a putative ligand-binding domain (LBD) classifies SHP as a member of the NR family, although the endogenous ligand has not been identified (Seol, Choi, & Moore, 1996) and NR0B2 is known as common heterodimer partner of FXR implicated in bile acid metabolism (Goodwin et al., 2000). Among best up-regulated genes found in PSC liver, FGF19 is implicate in bile acid homeostasis and effectively it is a target of bile acids via FXR in enterocyte and recirculate in portal blood to bind to FGFR4 receptor on hepatocyte (Inagaki et al., 2005). Among best up regulated molecules in liver of PSC, it was found TNFRSF12A which is a receptor from TNF superfamily. TNFRSF12A is also named TWEAK receptor and TWEAK/Fn14 signaling have been associated to promote cholangiocarcinoma niche formation and progression (Dwyer et al., 2021). Through liver functional analysis, FXR regulation dependency was used to build a liver protein-protein interaction network which mainly implicated immune response and nuclear receptor signalling. Through immune response functionality, CD274 (alias PDL1) was found regulated under liver FXR regulation. Effectively, some clinical trials in cancer therapy under penbrolizumab (anti-PDL1) highlighted risk to developed sclerosing cholangitis (Tahboub Amawi, Tremaine, & Venkatesh, 2020). In next step, human PSC gene signature was investigated with the FXR regulation dependency. NR0B2 was found as the best PSC predictive up regulated gene under FXR regulation and this relation is well known to be implicated in regulatory cascade of bile acid metabolism drive by nuclear receptors which repressed CYP7A1. CYP7A1 catalyzes the rate limiting step in bile acid biosynthesis. NR0B2 is known to repress CYP7A1 via activity inhibition of liver receptor homolog 1 (LRH-1) (Goodwin et al., 2000). In liver from PSC, CYP7A1 was confirmed to be down regulated as compared to liver samples from controls. Among best PSC predictive genes under FXR dependency, DGAT2, Diacylglycerol O-Acyltransferase 2 (Brandt, McFie, & Stone, 2016) implicated in biosynthesis of triglycerides was found repressed in PSC liver and effectively portal blood and bile lipidomics of PSC patients was found peculiar as compared to similar samples from disease controls and non-disease controls especially with increase of free fatty acid detection (Tietz-Bogert et al., 2018). NR0B2 alias SHP was found as the best predictive marker of PSC under FXR regulation dependency and its up regulation in PSC liver was found independent of the age, the gender and Body Mass Index (BMI) of the patients. Mdr2-/- alias Abcb4-/- mice model is widely used as a model of sclerosing cholangitis. It recapitulates the progressive fibrosing cholangitis aspect of human PSC. At single cell level in liver of Abcb4-/- mice, expression of Nr0b2 was detected in hepatocyte and cholangiocytes cell compartments but it’s over expression as compared to wide type cells was observed only the cholangiocytes which highly expressed Sox9. Based on alternative expression of Sox9 and Nr0b2, single cell trajectory was built with in Abcb4-/- cholangiocyte cell compartment. Affected cholangiocytes highlighted a cell trajectory cluster comprising a network of molecules implicated in metabolism: sulfur component metabolism, glutathione derivative biosynthesis and monocarboxylic acid metabolism. N-acetylcysteine therapeutical option have been already tested in PSC, by its antimucolytic action it potentially reduce bile viscosity and improved clinical and laboratory parameters of PSC patients in a small study (Ozdil, Cosar, Akkiz, Sandikci, & Kece, 2011). On Nr0b2 cell trajectory of affected cholangiocyte, Gsta3 Glutathione S transferase was found with similar pseudotime expression meaning their closed regulation during cell trajectory of this affected cell compartment. Glutathione S-transferases (GST) like GSTA3 acts as antioxidant enzymes mainly in liver tissue through glutathione peroxidase activity toward phospholipid hydroperoxides (Yang, Sharma, Zimniak, & Awasthi, 2002). GSTA3 belongs to the GST α-class, which converts lipid peroxides to glutathione conjugates (Johansson & Mannervik, 2001, p. 3-3). It has been shown that glutathione S-transferase A3 have a role in inhibiting hepatic stellate cell activation and rat hepatic fibrosis (H. Chen et al., 2019). Also, glutathione S-transferase A3 knockout mice harbores liver injury, oval cell proliferation and cholangiocarcinogenesis in (Crawford et al., 2017). Inhibitor of DNA binding 2 (ID2) was found also closely regulated on Nr0b2 cell trajectory of Abcb4-/- cholangiocytes. ID2 is a member basic Helix Loop Helix transcription factor family. ID2 like other ID proteins lacks DNA binding domain and so transcription of downstream cell cycle targets like p16, p21(Benezra, Davis, Lockshon, Turner, & Weintraub, 1990). ID2 like ID1 and ID3 proteins is frequently overexpressed in all biliary tract cancer (BTC) subtypes. Also, ID protein expression is deregulated in many tumors including pancreatic cancer a malignancy somehow related to bile tract cancers. During bile tract cancer, for subgroup of patients who had received chemotherapy, if nuclear expression of ID2 is negative these patients presented a better overall survival as compared those who were nuclear positive (Harder et al., 2013). With closed pseudotime expression on Nr0b2 cell trajectory of Abcb4-/- cholangiocyte cell compartment, it was found the membrane molecule Tmem45a. Tmem45a is transmembrane molecule of 275 amino acids, predicted to have five to seven transmembrane domains and localized in the trans Golgi apparatus. This protein is over expressed in many cancers: breast cancer, liver cancer, renal cancer, glioma, head and neck cancer, ductal cancer and ovarian cancer: it could be supposed to act as an oncogene (Flamant et al., 2012; Guo et al., 2015; Lee et al., 2012; Manawapat-Klopfer et al., 2016; Sun et al., 2015; Wrzesiński et al., 2015). During breast and cervical tumors, there was found a higher level of expression for TMEM45A suggesting this molecule could be a biomarker of aggressiveness in these malignant pathologies (Manawapat-Klopfer et al., 2016). TMEM45A have been implicated in the proliferation, migration and invasion of glioma cancer cells (Guo et al., 2015; Sun et al., 2015). TMEM45A also confers chemoresistance in hypoxia condition for the cell death induced chemotherapeutic agents of breast cancer (Flamant et al., 2012).

## Conclusion

In this work we performed integration of biomedical text-mining related to PSC symptom in omics data such as human transcriptome of PSC liver, FXR functional liver transcriptome and liver single cell transcriptome of the Abcb4-/- model of PSC. This integrative work allowed to confirmed major implication of Nr0b2 and its associated nuclear receptors like FXR in a metabolic cascade that could influence immune response. TNFRSF12A/TWEAK receptor, was found up regulated in PSC liver independently of FXR regulation and TWEAK signaling is known for its implication in pre-conditioning niche of cholangiocarcinoma. At single cell level, Nr0b2 up regulation was found in cholangiocytes but not in hepatocytes. In this affected cell compartment cell trajectory defined on Nr0b2 expression, highlighted several metabolic pathways of detoxification like sulfur, glutathione derivative and monocarboxylic acid metabolisms. On this cell trajectory it was discovered some molecules potentially implicated in carcinogenesis like: GSTA3, ID2 and mainly TMEM45A a transmembrane molecule considered as oncogene in several cancers.

## DATA AVAILABILITY

R bioinformatics code for single cell analyses performed with Seurat 4 and Monocle 2 packages on dataset GSE168758 are accessible on internet at the following address: www.github.com/cdesterke/pscsc (accessed on 2021, July 10^th^).

## ACKNOWLEDGMENTS

Thanks to department ‘Bloc OPératoire Augmenté’ (BOPA) for access to their facilities.

## CONTRIBUIONS

CD and CF : design of the experience, CD and CF : written manuscript, CD : data analysis, CF : suppervisor and correction of the manuscript

## ABBREVIETIONS

Abcb4: ATP Binding Cassette Subfamily B Member 4
BMI: body mass index
BTC: biliary tract cancer
DEGs: differential expressed genes
FDR: False Discovery Rate
FXR: Farnesoid X Receptor, alias NR1H4
GO-BP: Gene Ontology Biological Process database
GSEA: Gene Set Enrichment Analysis
GW4064: Farnesoid X Receptor agonist
ID2: inhibitor of DNA binding 2
KO: knock-out
LIMMA: LInear Model from Microarray
MeSH: Medical Subject Headings
NAFLD: Nonalcoholic fatty liver disease
NASH: Nonalcoholic steatohepatitis
NR: Nuclear receptor
Nr0b2: Nuclear Receptor Subfamily 0 Group B Member 2, alias
SHP: Small Heterodimer Partner
PBC: primary biliary cholangitis
PPI: Protein-protein interaction
PSC: primary sclerosing cholangitis
PTM: Pavlidis Template Matching
t-SNE: t-distributed stochastic neighbor embedding
UMAP: Uniform Manifold Approximation and Projection

## SUPPLEMENTAL MATERIAL

### Supplemental Figures

**Supplemental Figure 1:**
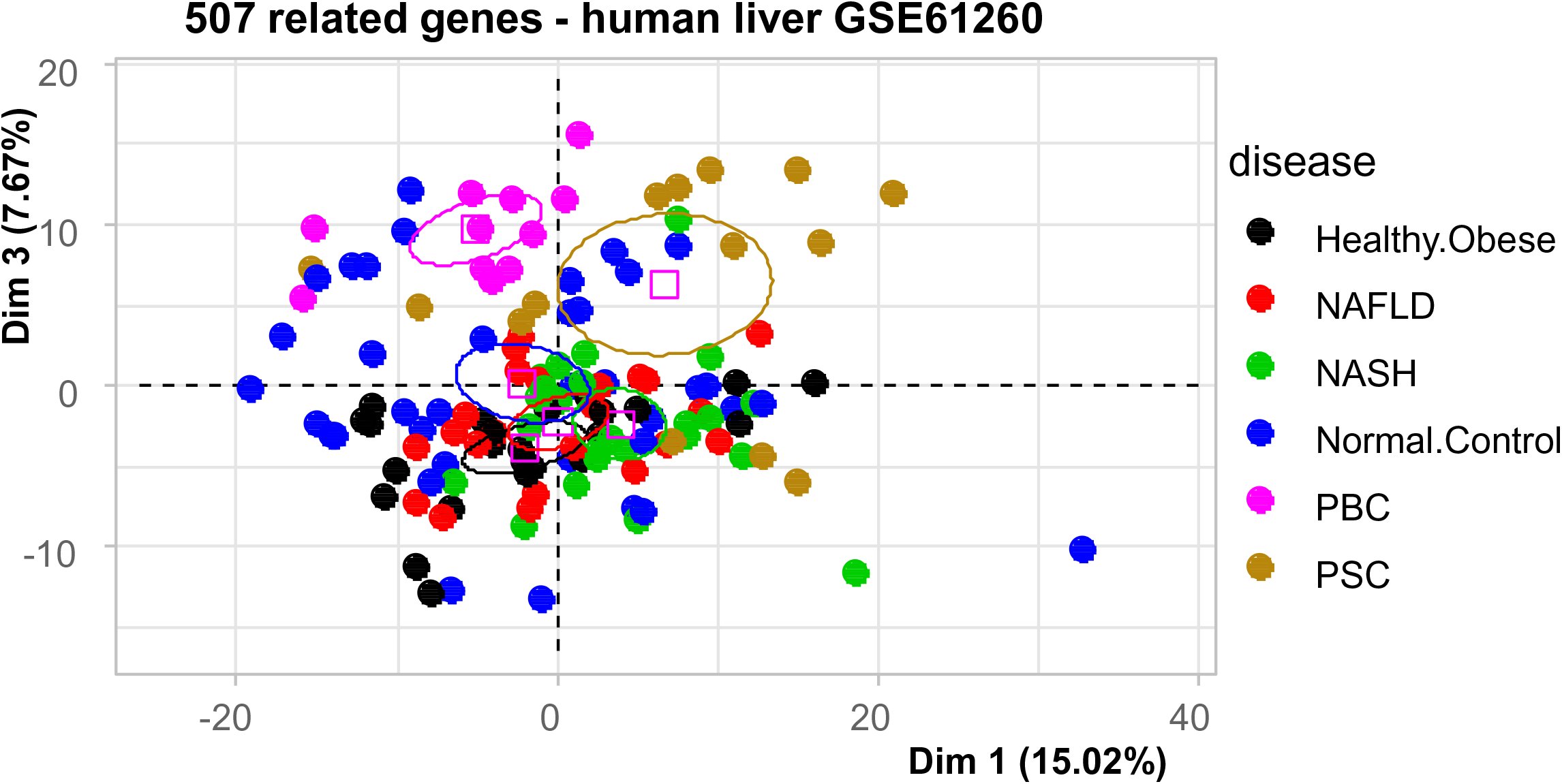
Stratification of PSC and PBC liver transcriptomes by unsupervised analysis: Second principal map of principal component analysis performed on liver transcriptomes of dataset GSE61260 with PCS related genes found by text mining.

**Supplemental Figure 2:**
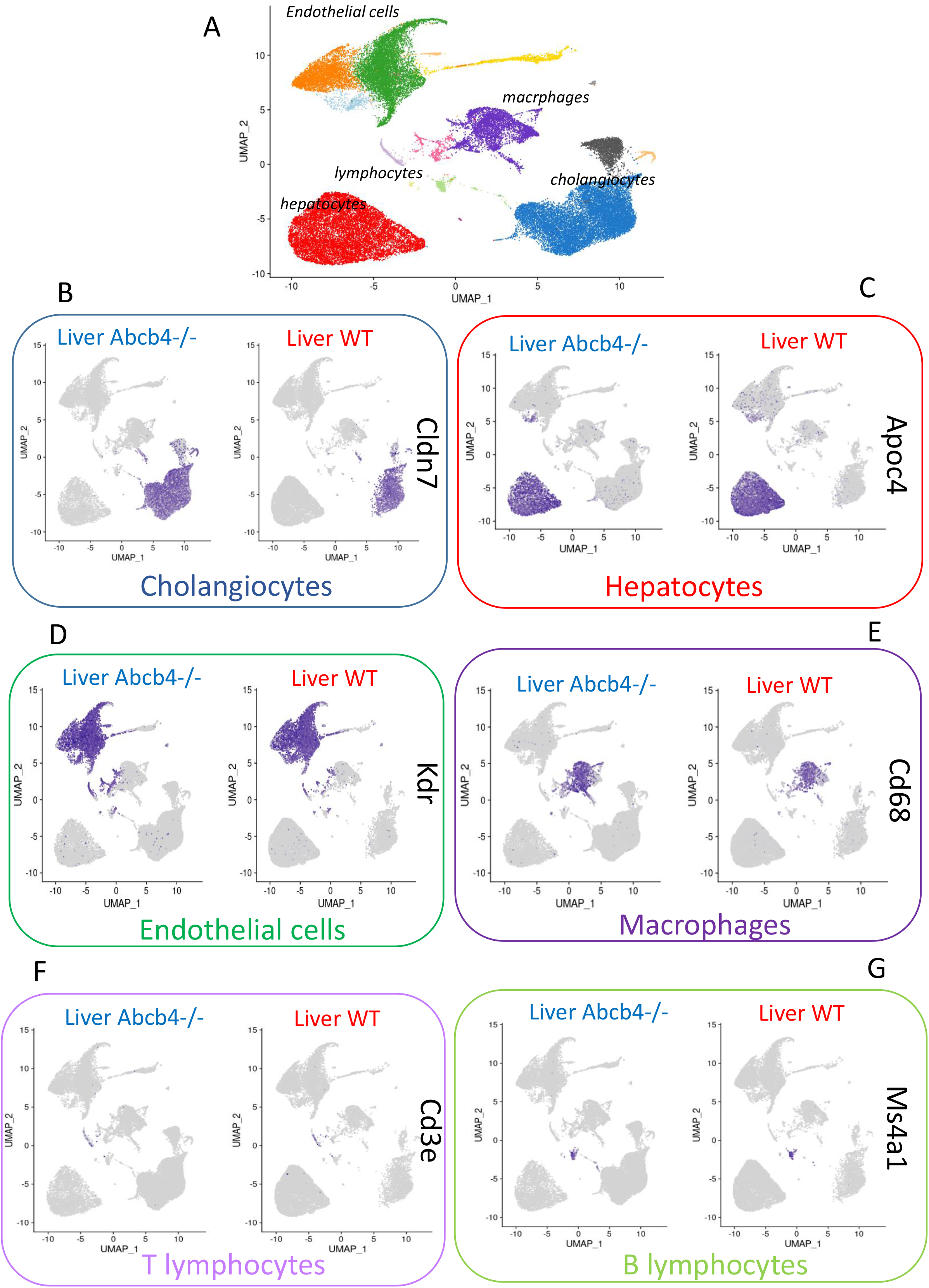
Single cell heterogeneity of clusters identified in WT and Abcbc4-/- livers: A/ whole liver UMAP dimension reduction with summary of the cell cluster indentifications; B/ Cldn7 featureplot which identified cholangiocyte cell subgroup; C/ Apoc4 featureplot which identified hepatocyte cell subgroup; D/ Kdr featureplot which identified endothelial cell subgroup; E/ Cd68 featureplot which identified macrophage cell subgroup; F/ Cd8a featureplot which identified T-lymphocyte cell subgroup; G/ Ms4a1 Kdr featureplot which identified B-lymphocyte cell subgroup.

**Supplemental Figure 3:**
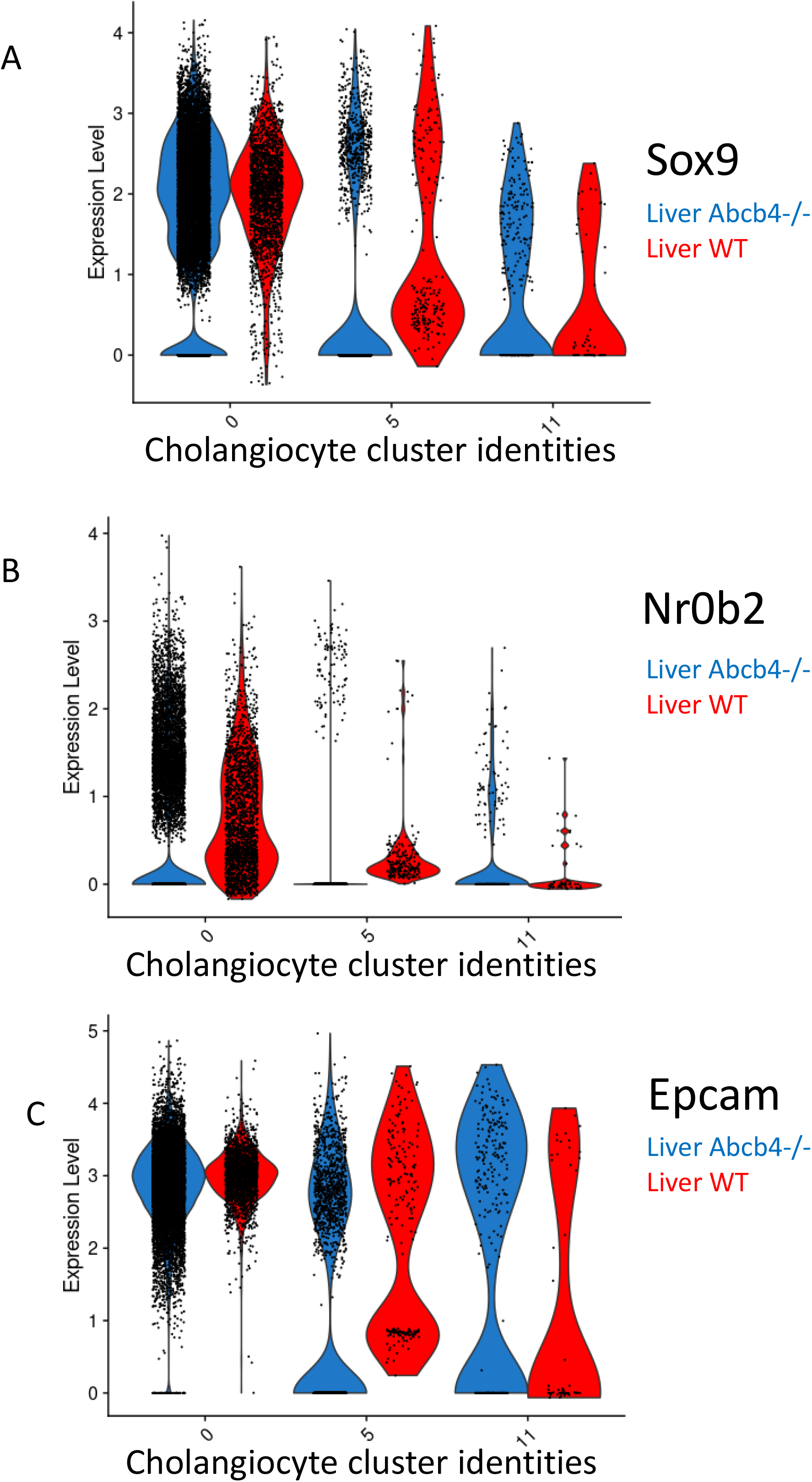
Single cell differential expression of Sox9, Nr0b2, Epcam in cholangiocyte clusters of WT and Abcb4-/- livers: A/ Violinplot of Sox9 expression in cholangiocyte cluters 0-5-11 stratified on phenotype wildtype versus Abcb4-/-; B/ Violinplot of Nr0b2 expression in cholangiocyte cluters 0-5-11 stratified on phenotype wildtype versus Abcb4-/-; C/ Violinplot of Epcam expression in cholangiocyte cluters 0-5-11 stratified on phenotype wildtype versus Abcb4-/-;

**Supplemental Figure 4:**
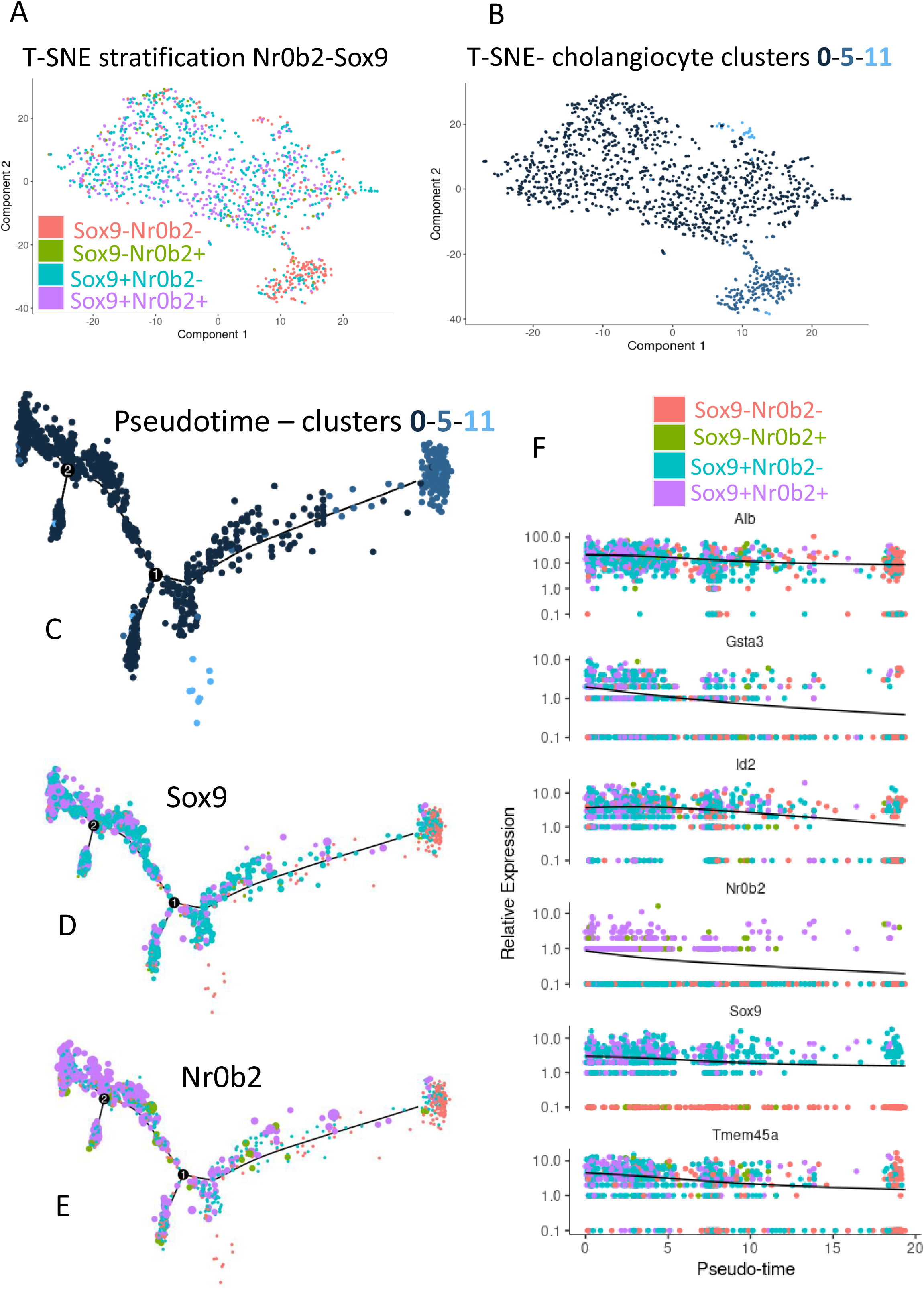
Pseudotime transformation of cholangiocyte cell trajectory in Abcb4 -/- livers: A/t-SNE plot of Abcb4-/- cholangiocyte with group stratification based on alternative expression of Sox9 and Nr0b2; B/ t-SNE plot of Abcb4-/- cholangiocyte with group stratification identified in Seurat clustering: clusters 0-5-11; C/ pseudotime tree with group stratification identified in Seurat clustering: clusters 0-5-11; D/ pseudotime tree with expression of sox9 as dot size and group stratification as dot color; E/ pseudotime tree with expression of Nr0b2 as dot size and group stratification as dot color; F/ pseudotime expression plot for markers found closely regulated with Nr0b2 on pseudotime cell trajectory

### Supplemental Tables

**Supplemental Table 1:**
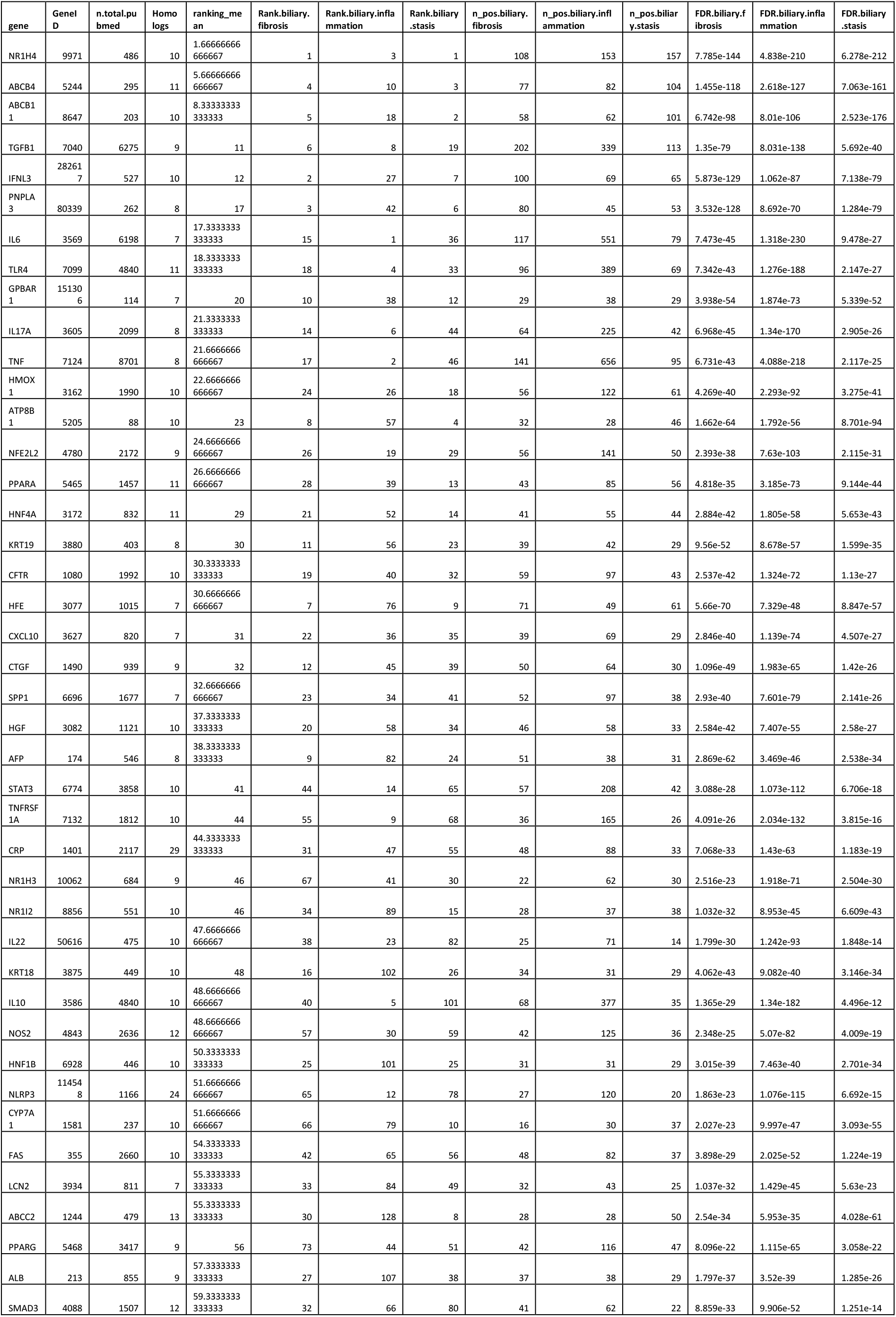
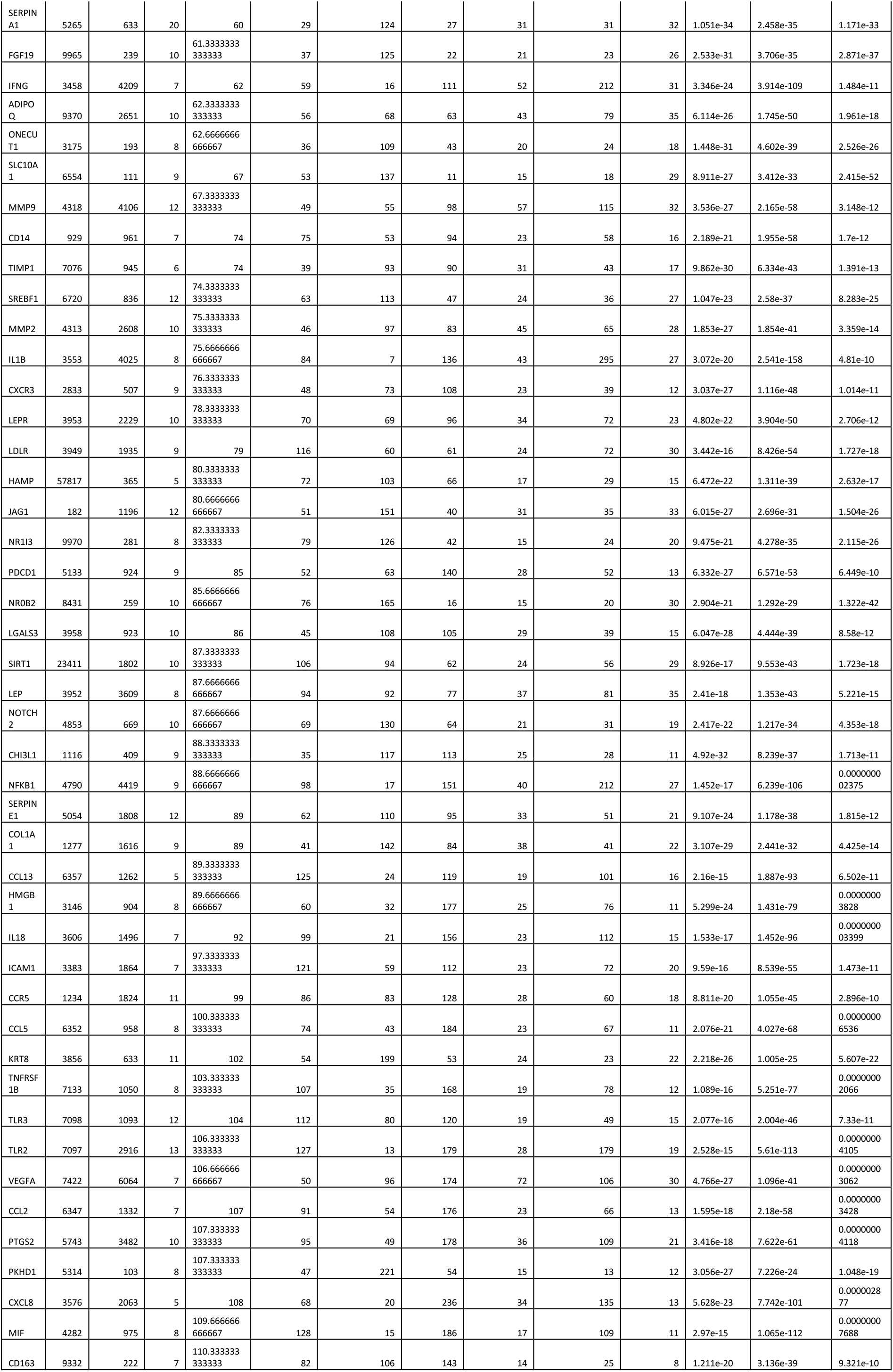
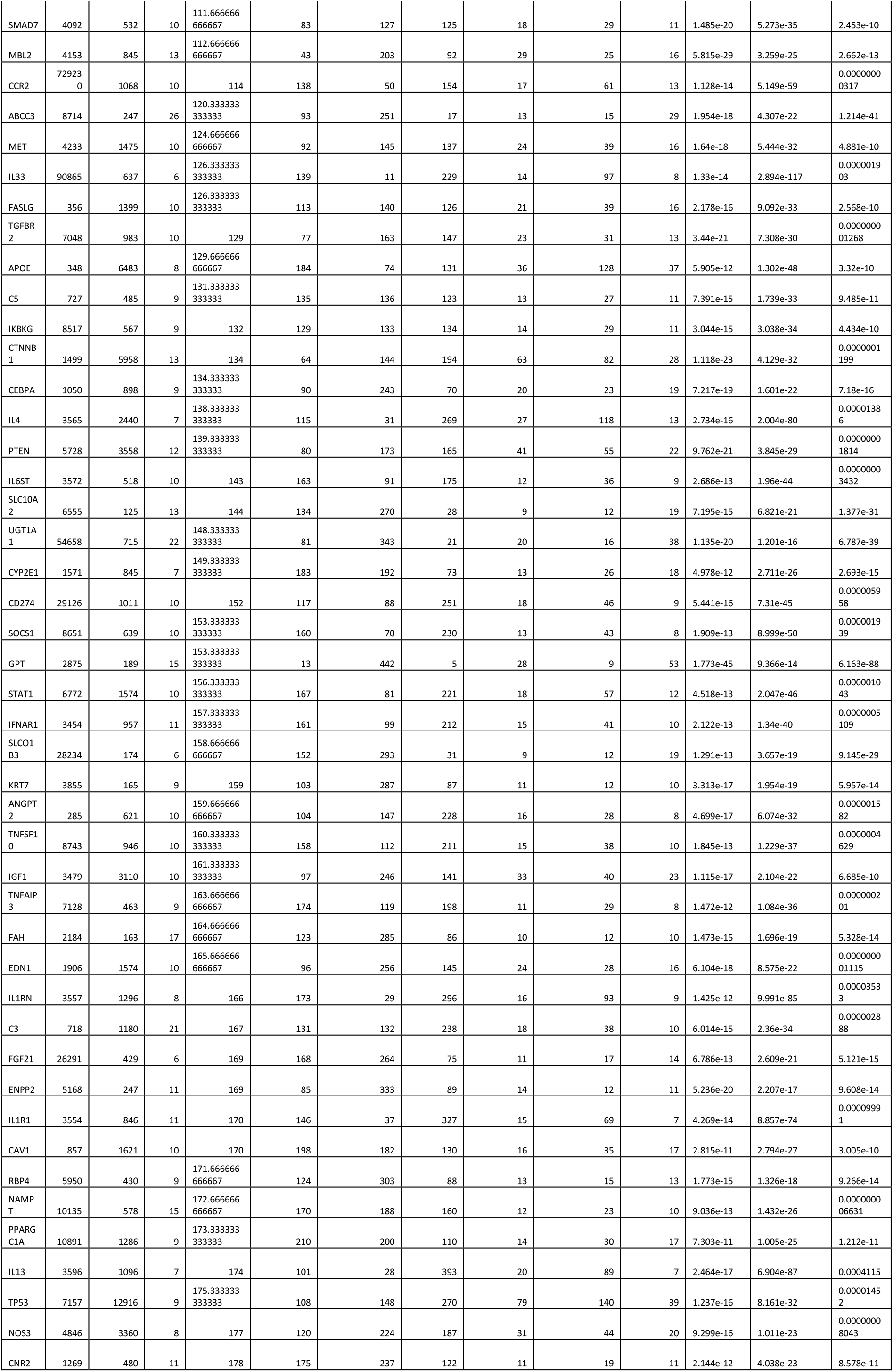
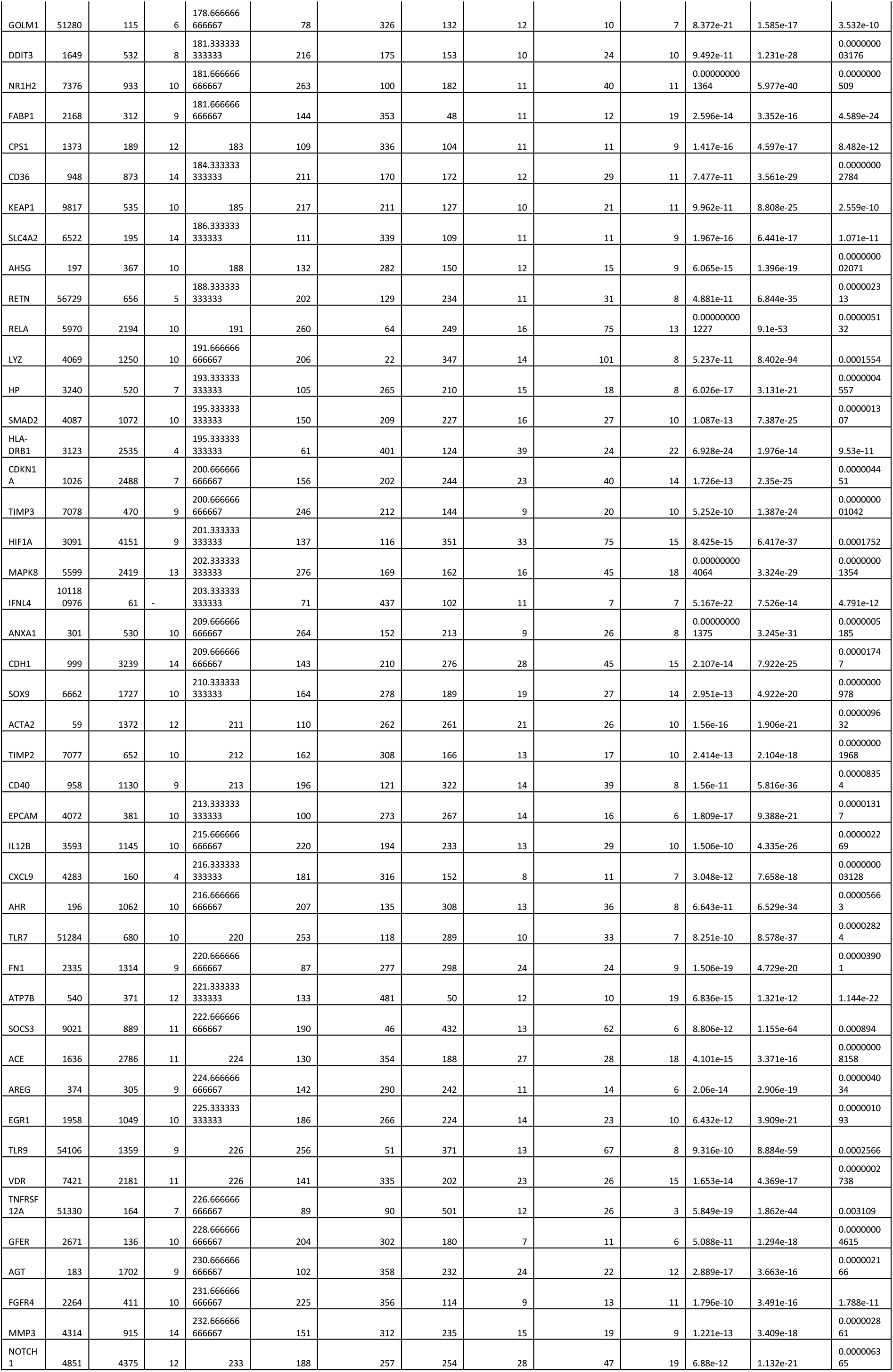
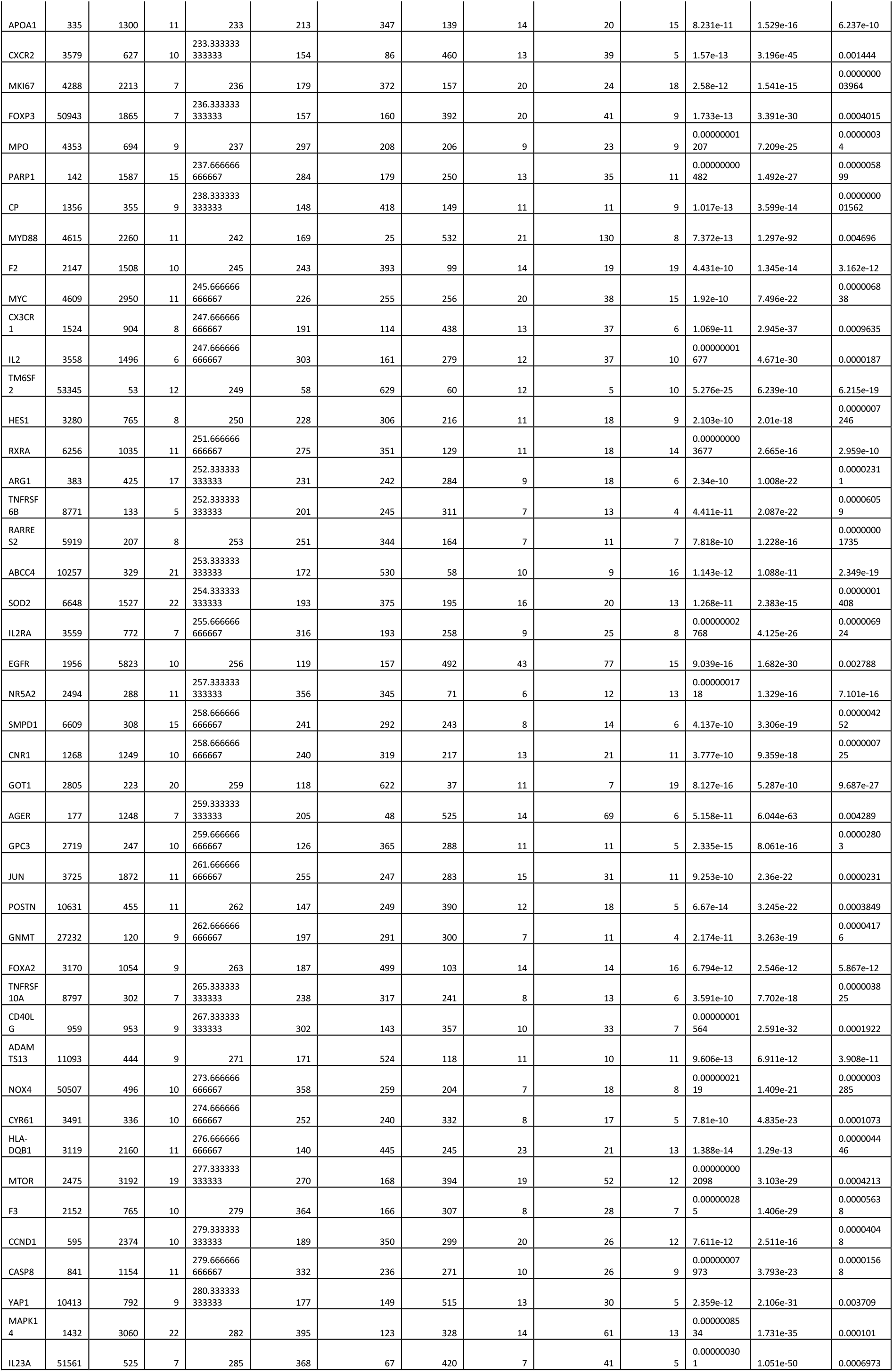
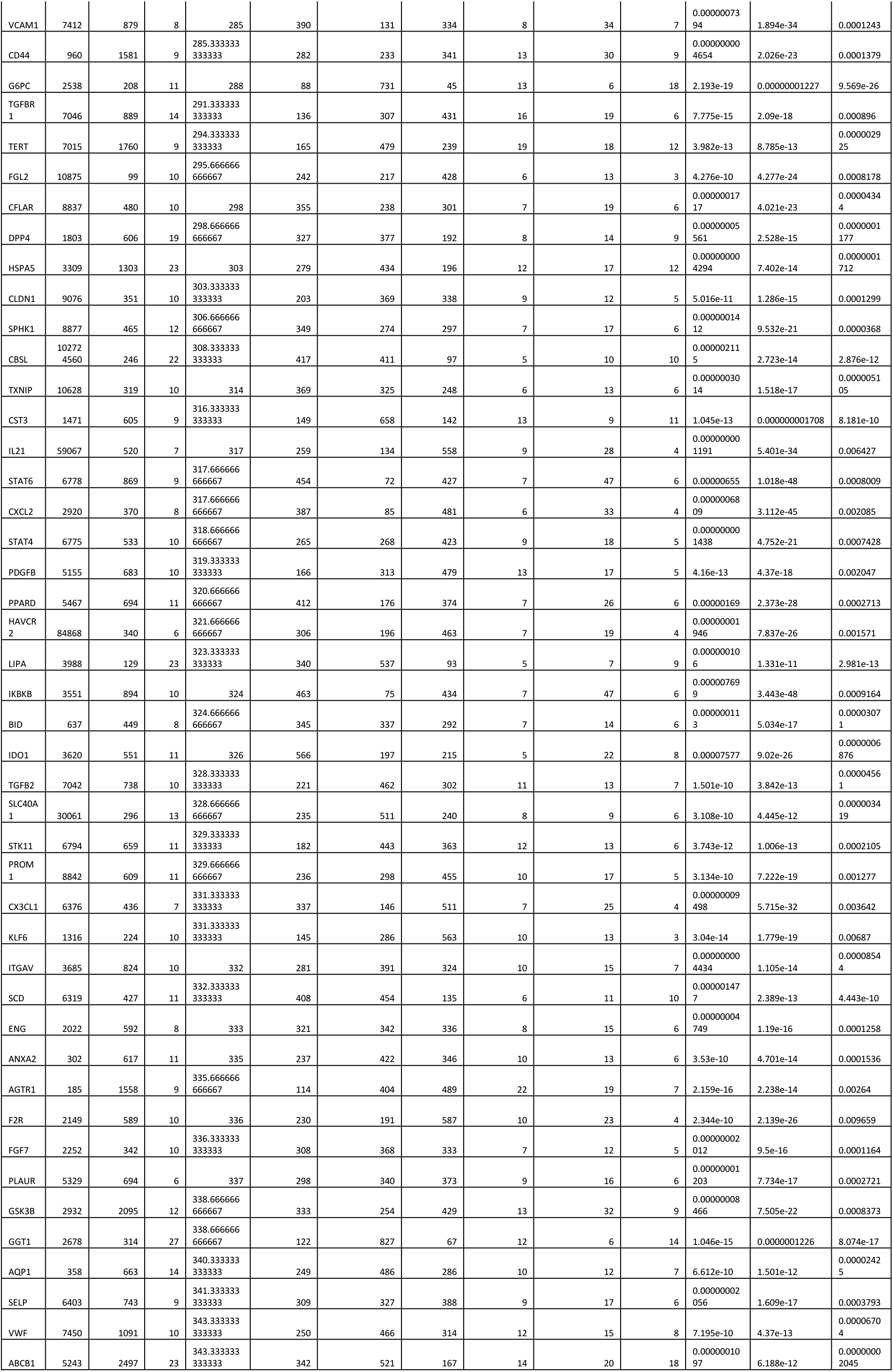
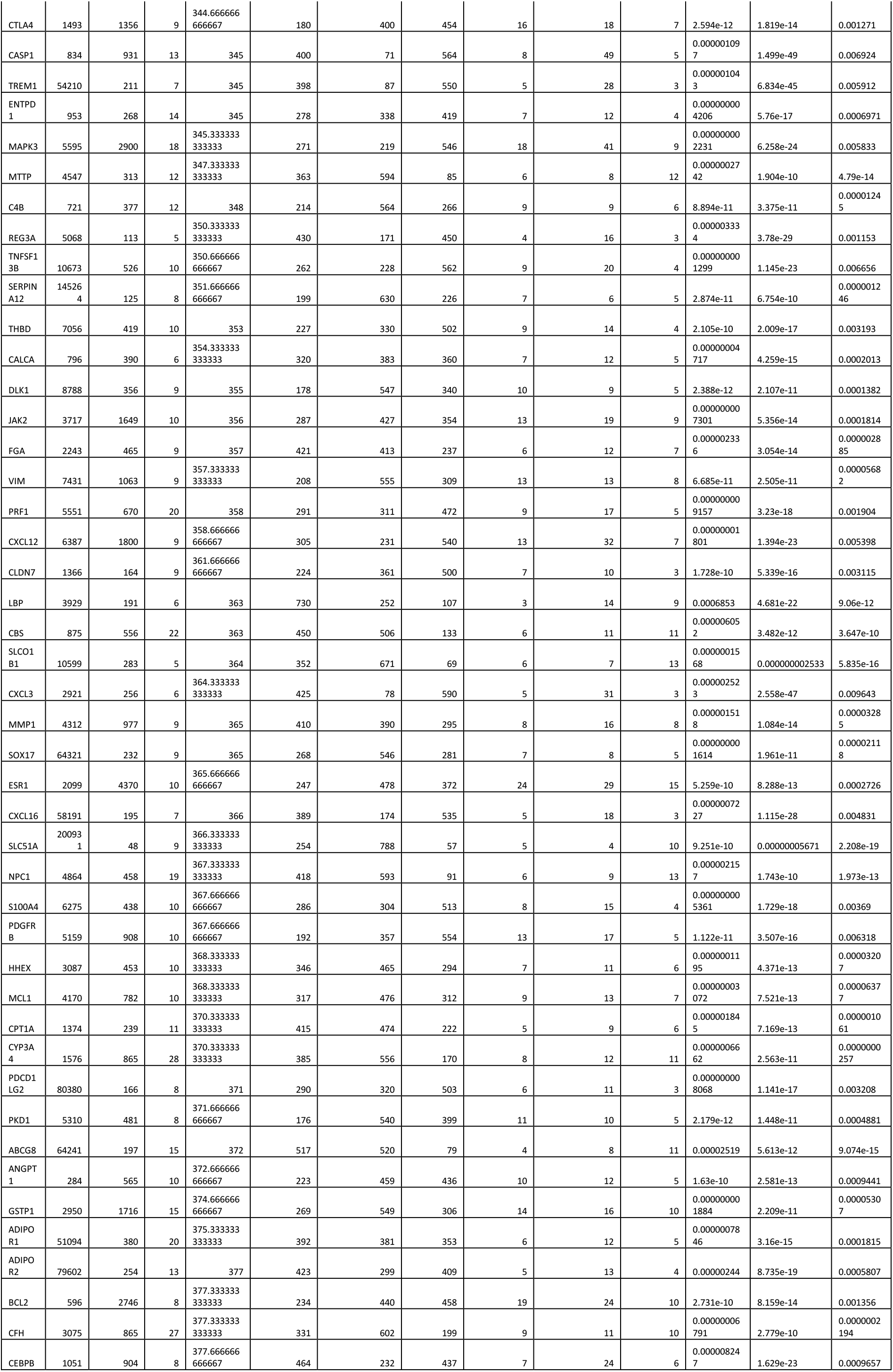
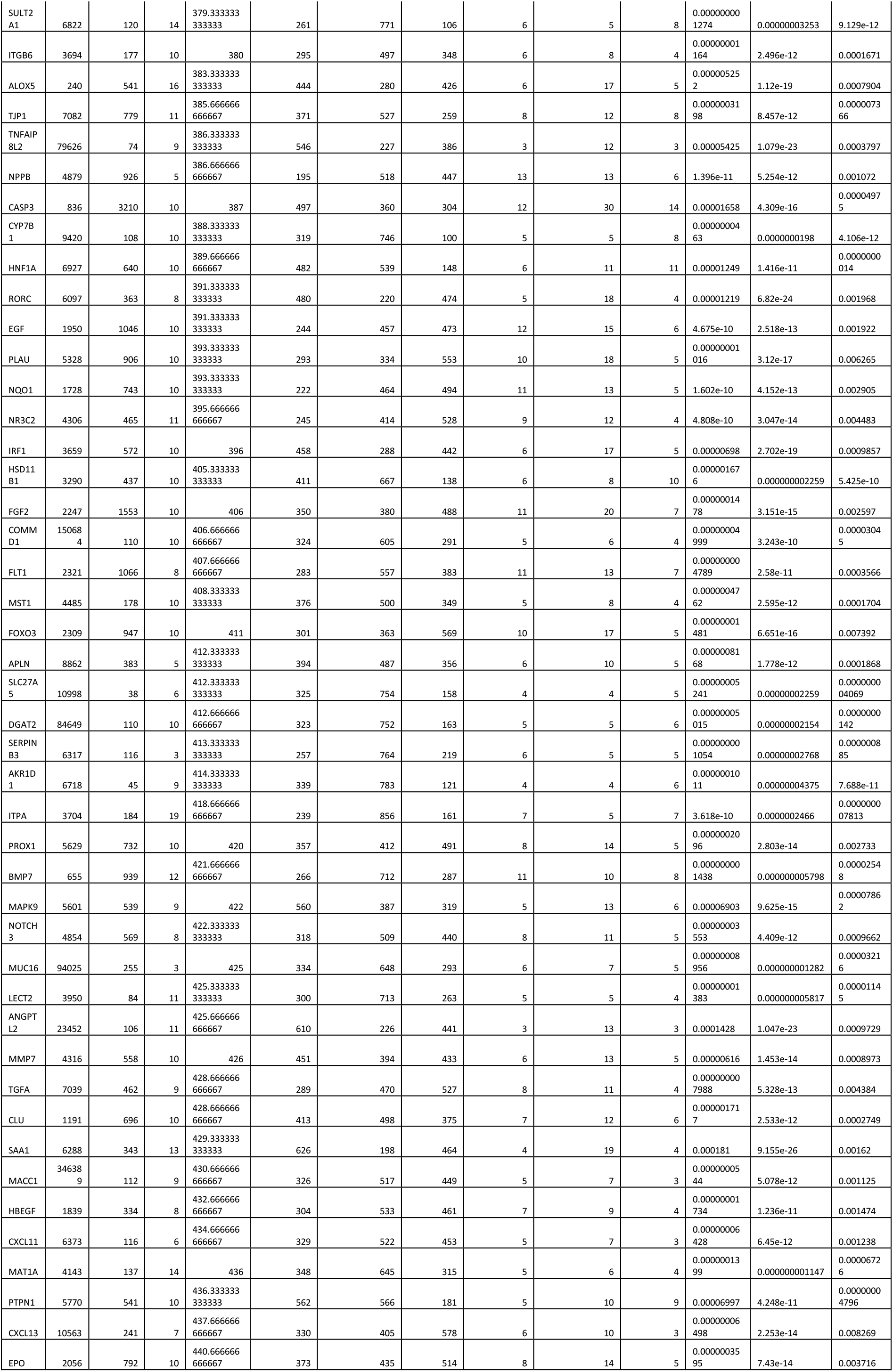
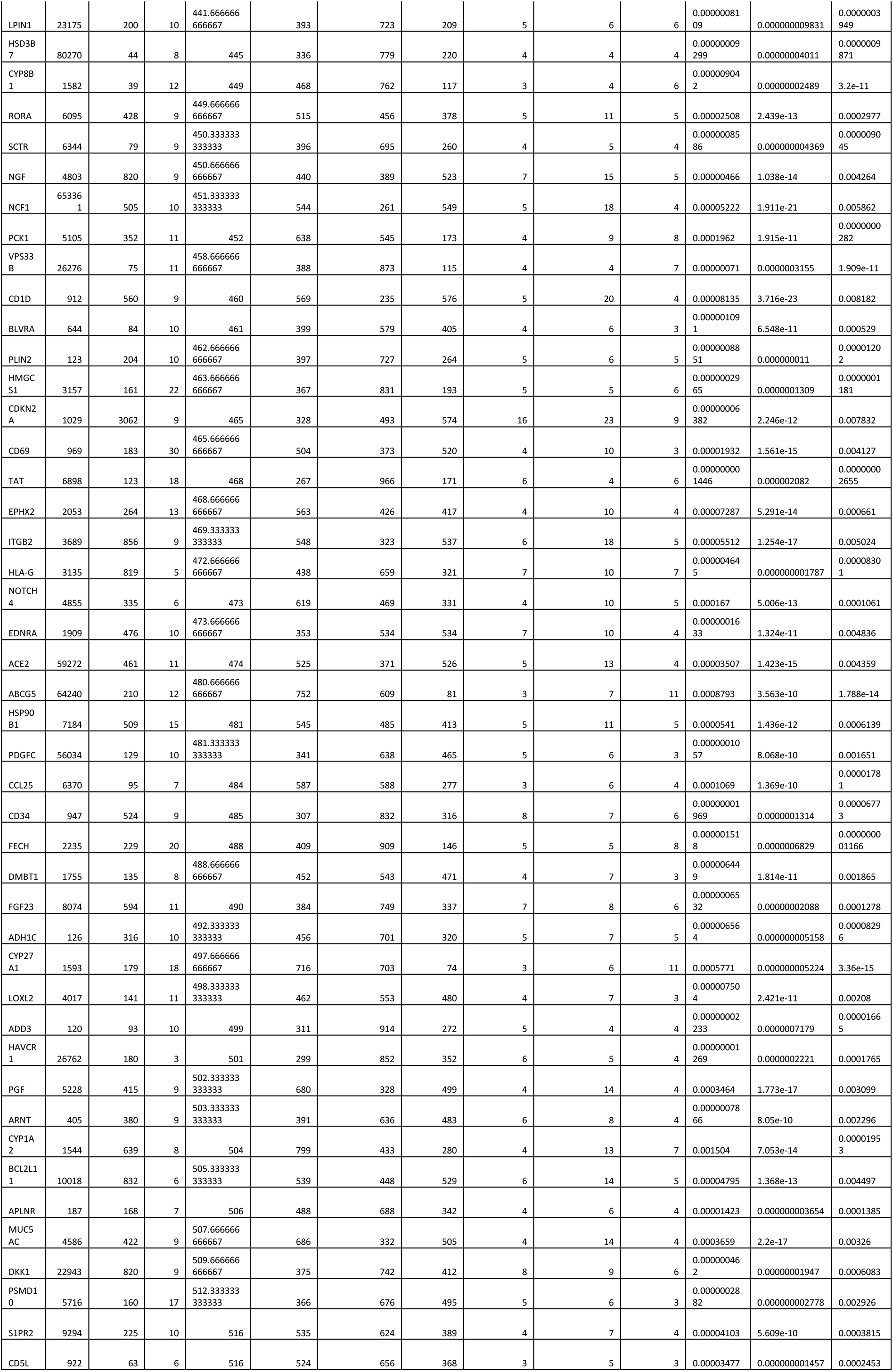
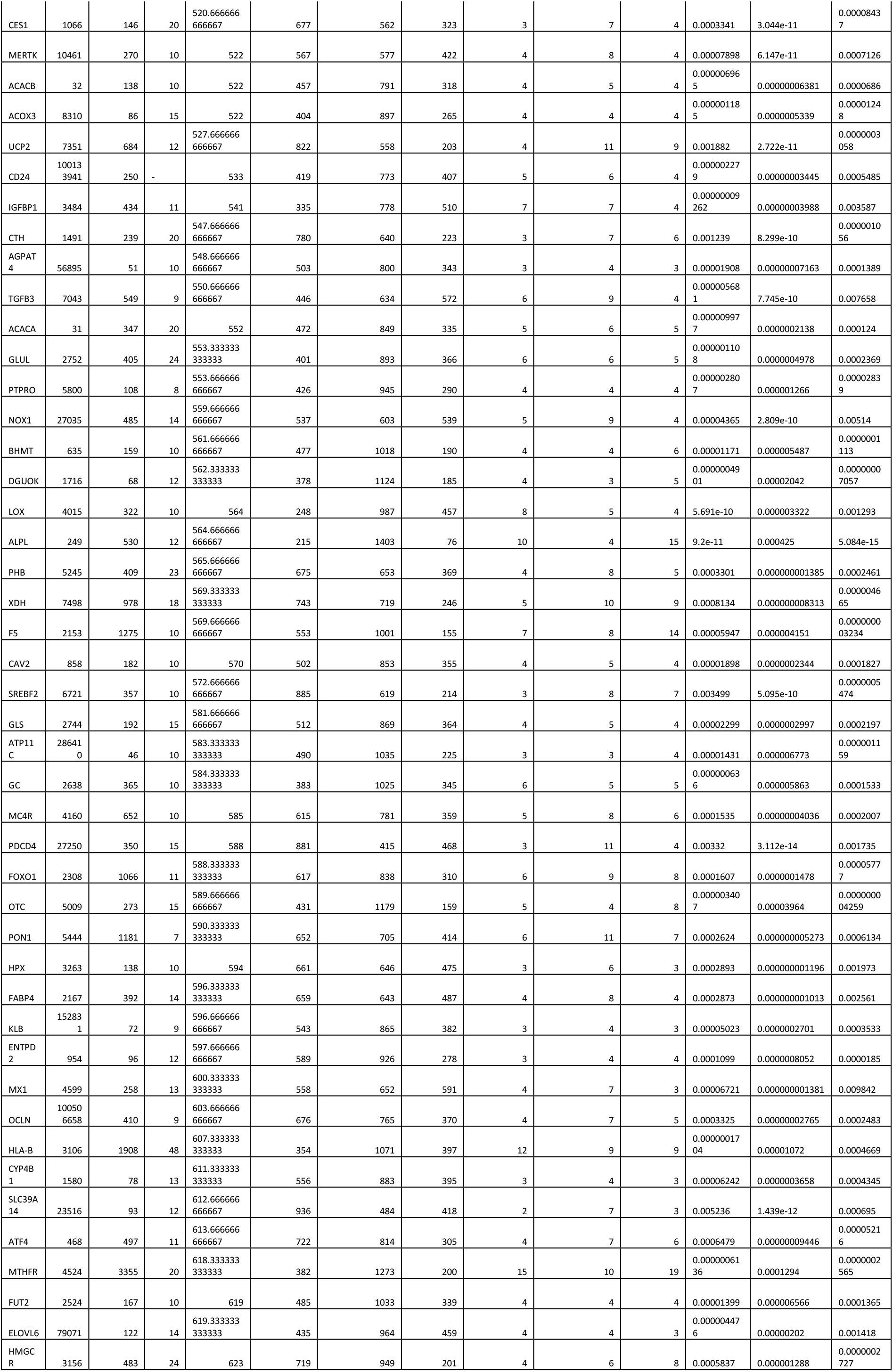
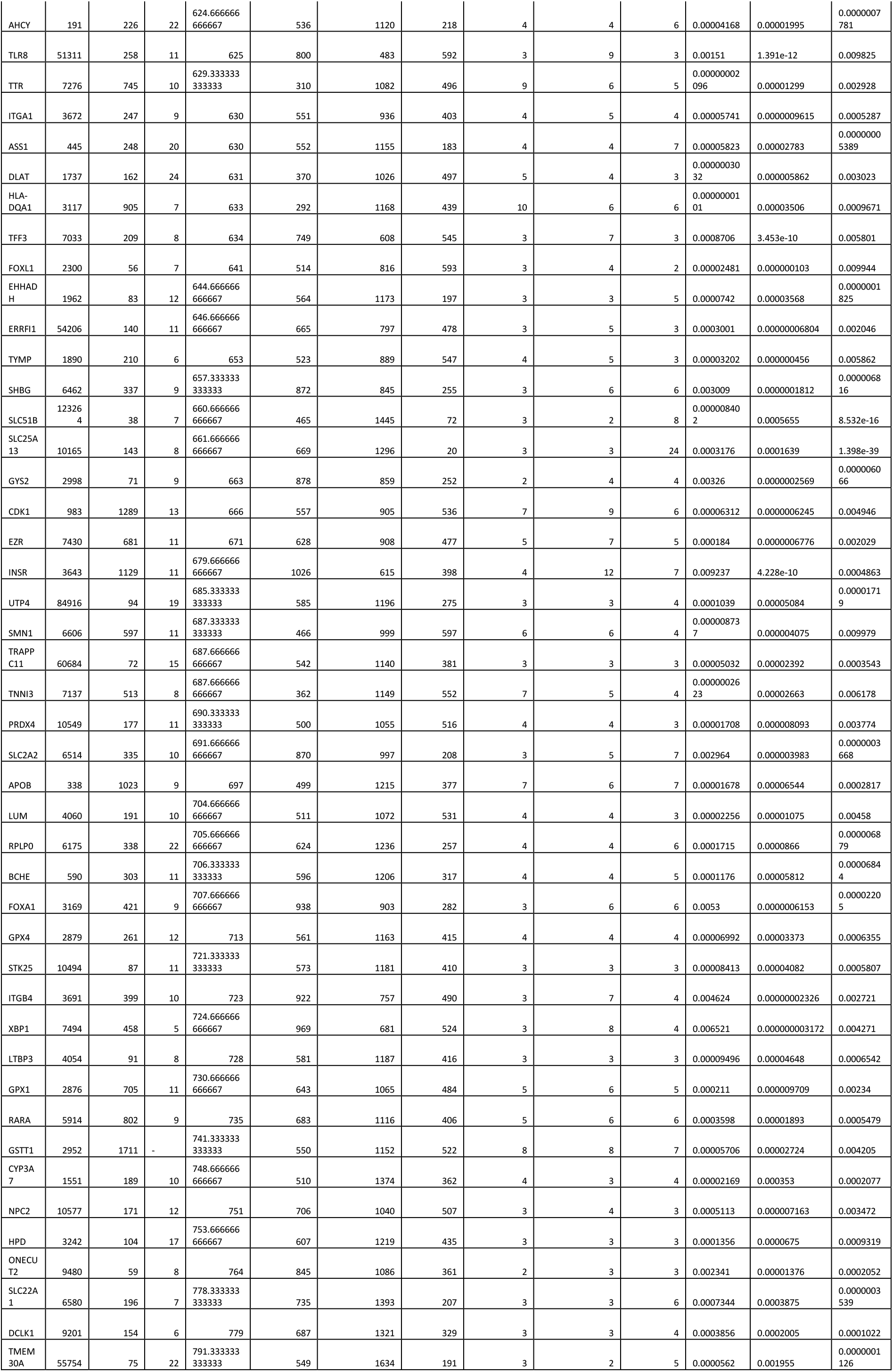
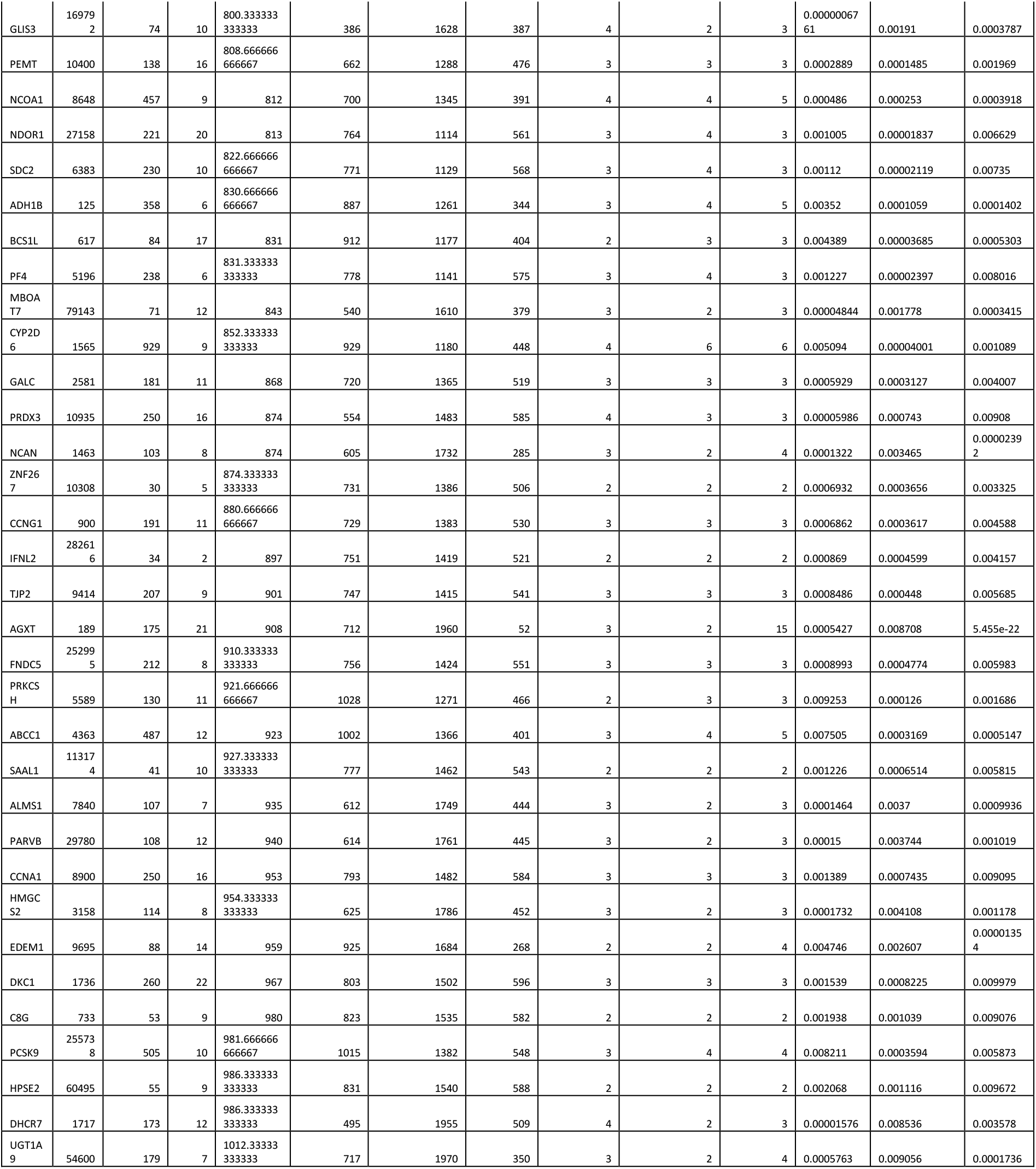
**Textmining table of the common 525 ranked genes found in Pubmed with symptom keywords:** this table describes the 525 ranked genes obtained by text mining with ‘Génie’ algorithm against the three MESH terms: biliary inflammation, biliary fibrosis, biliary stasis, respective ranks, p-values and collected PMID identifiers for each gene and each MESH term are presented

**Supplemental Table 2:**
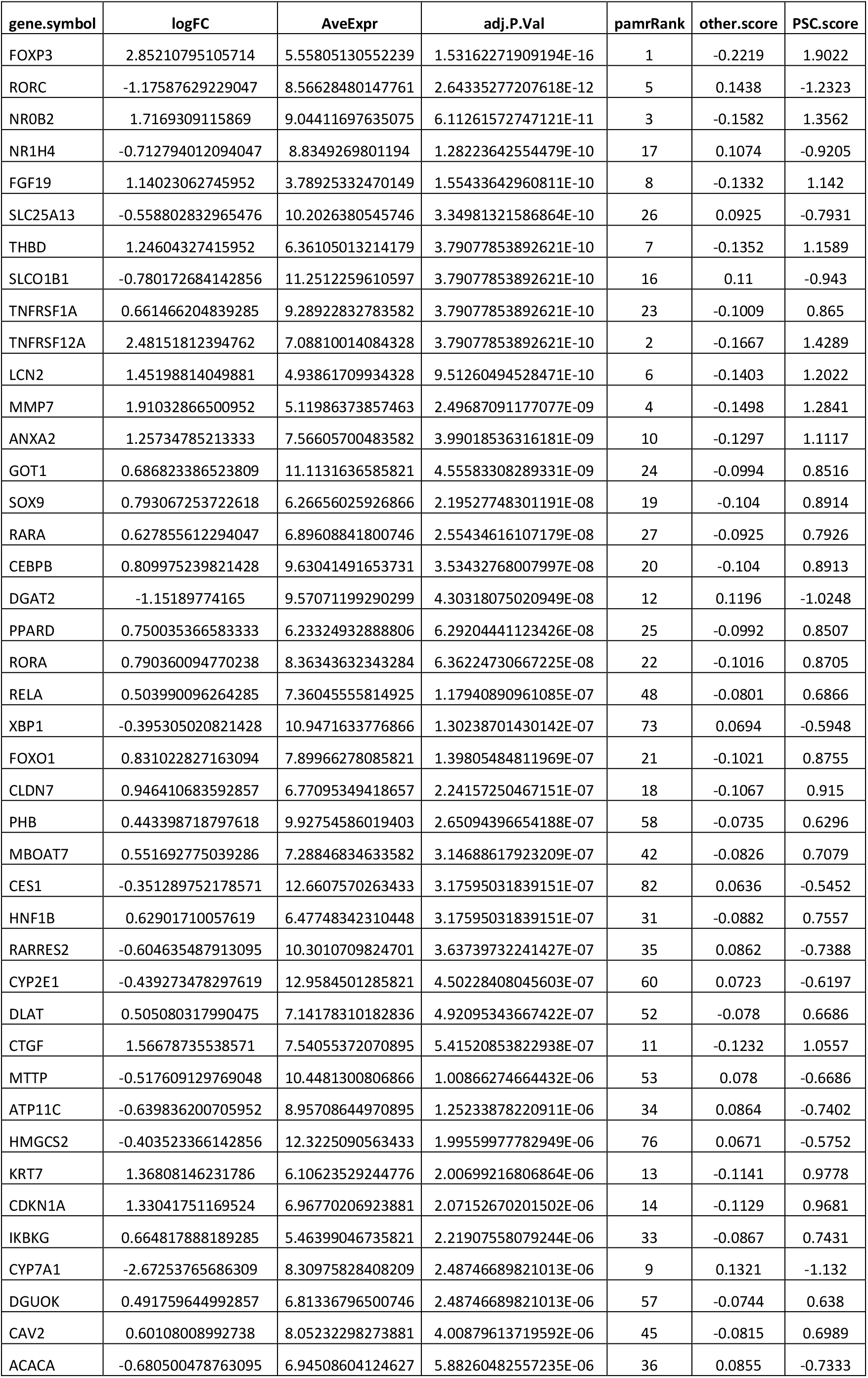
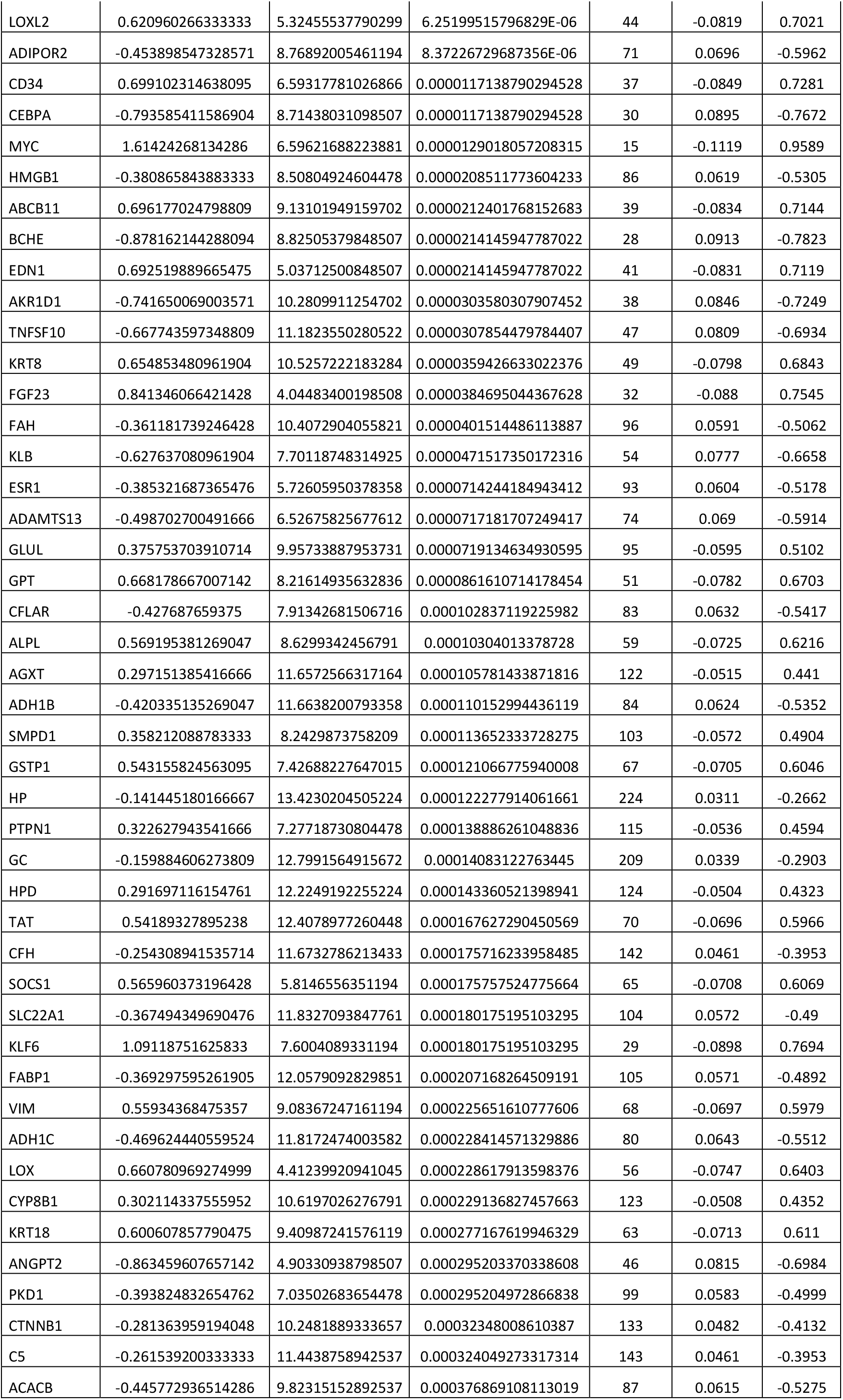
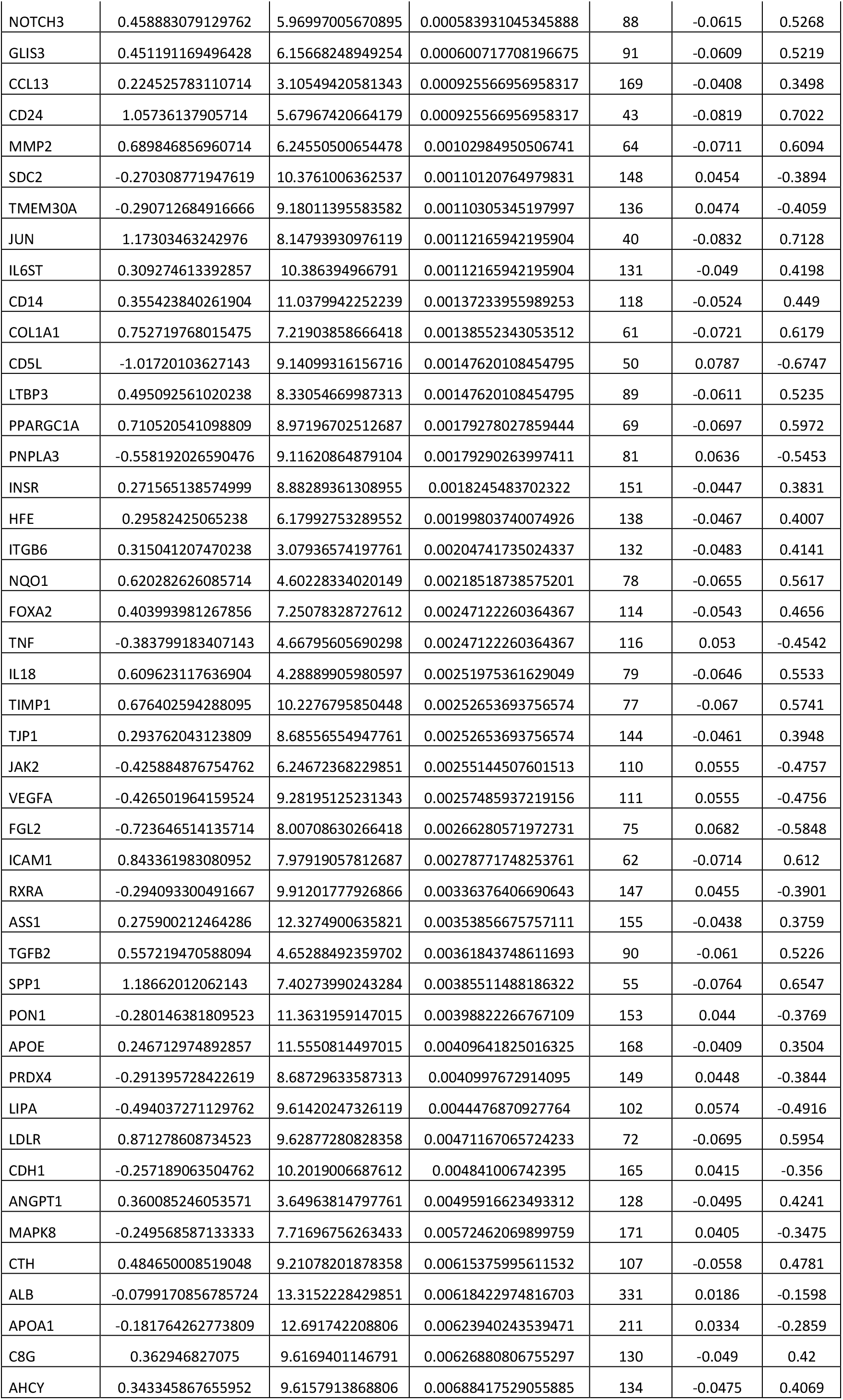
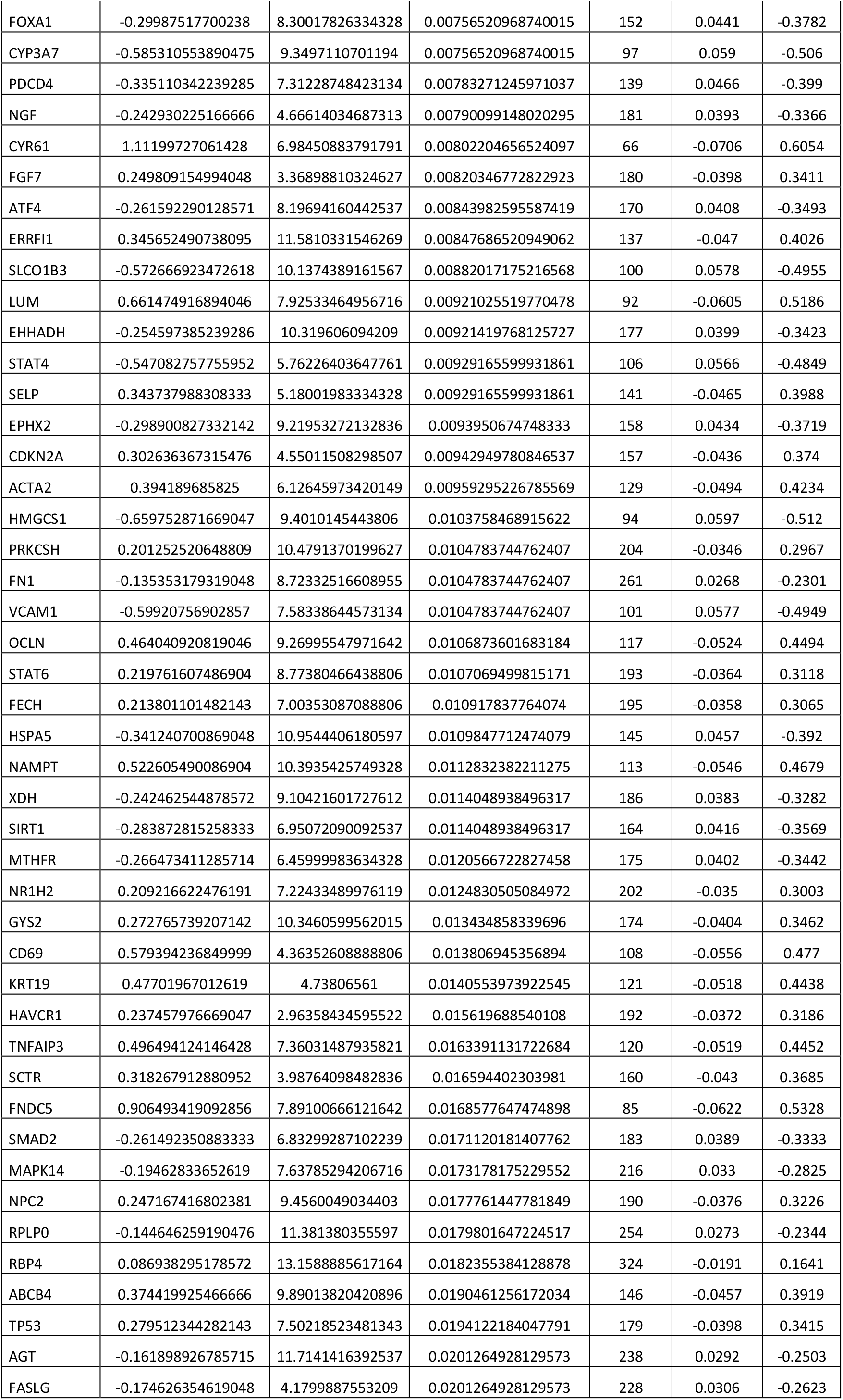
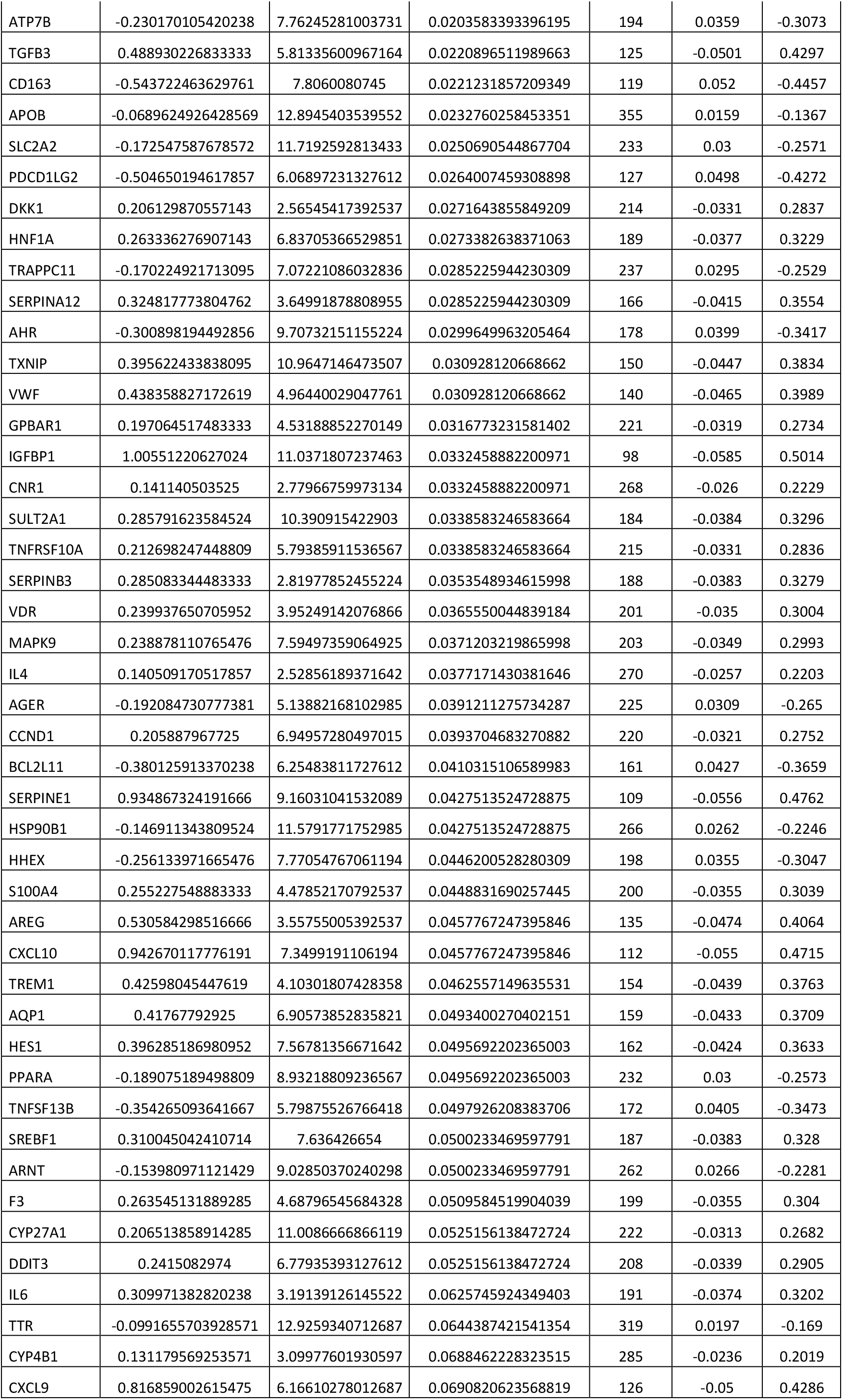
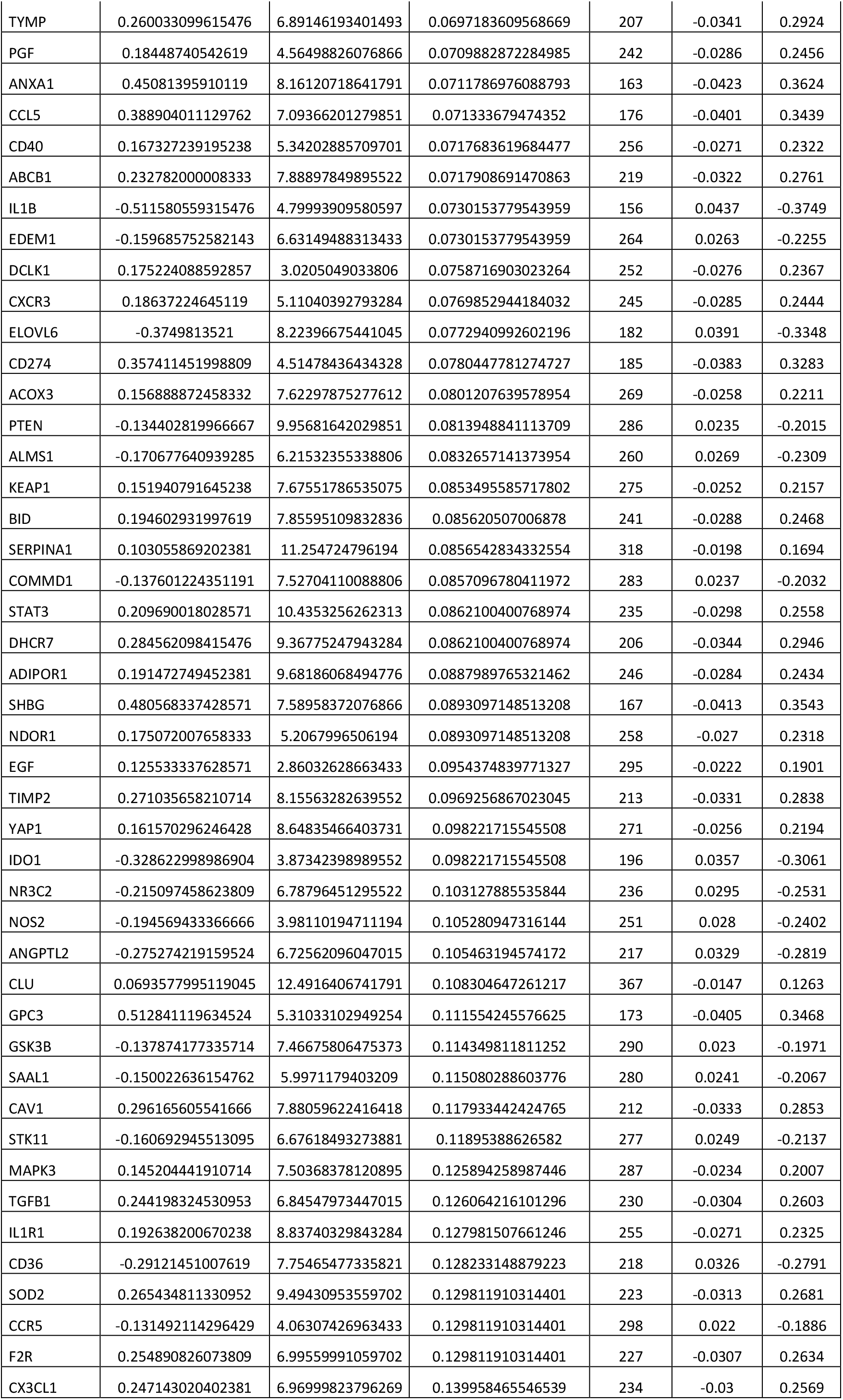
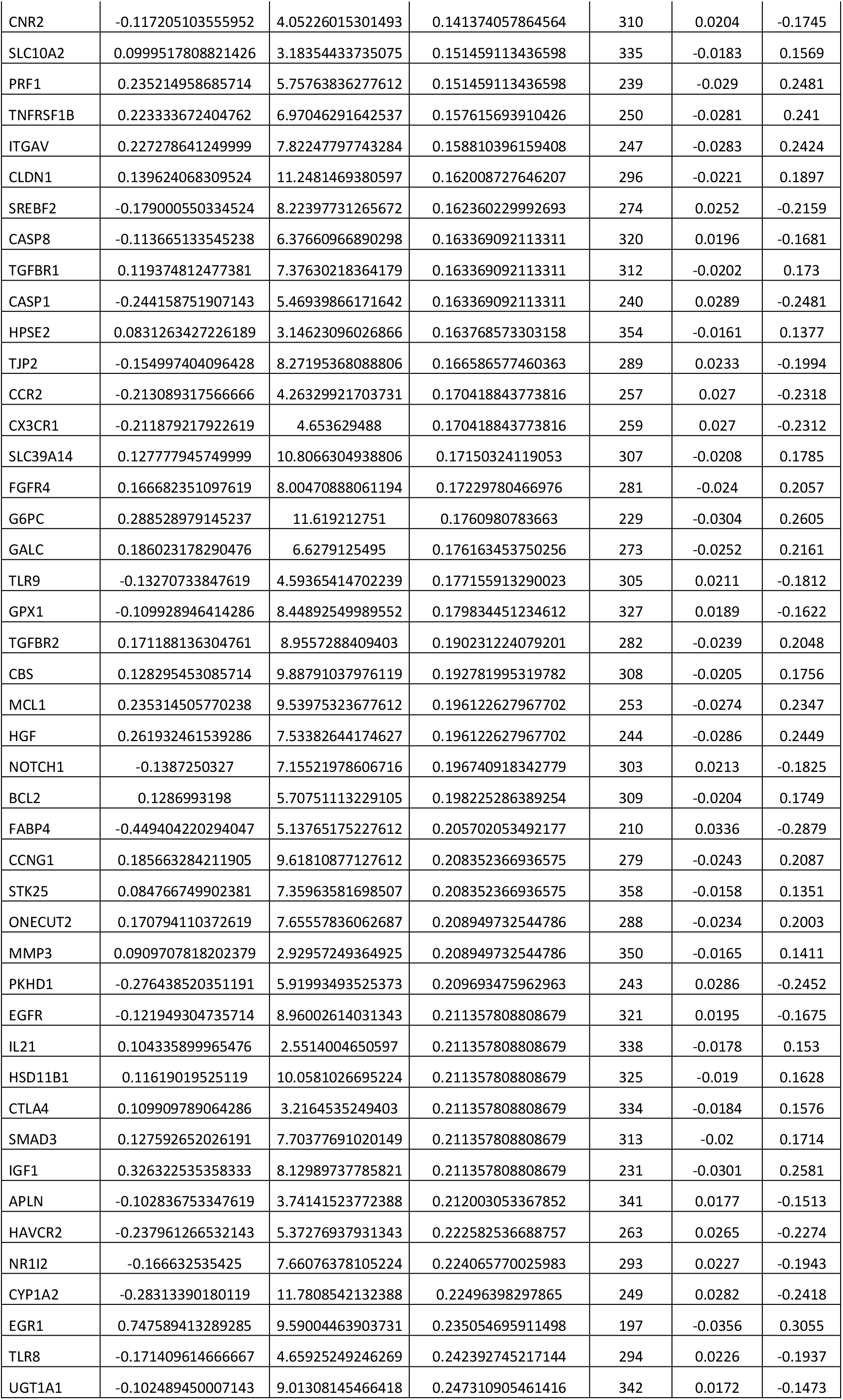
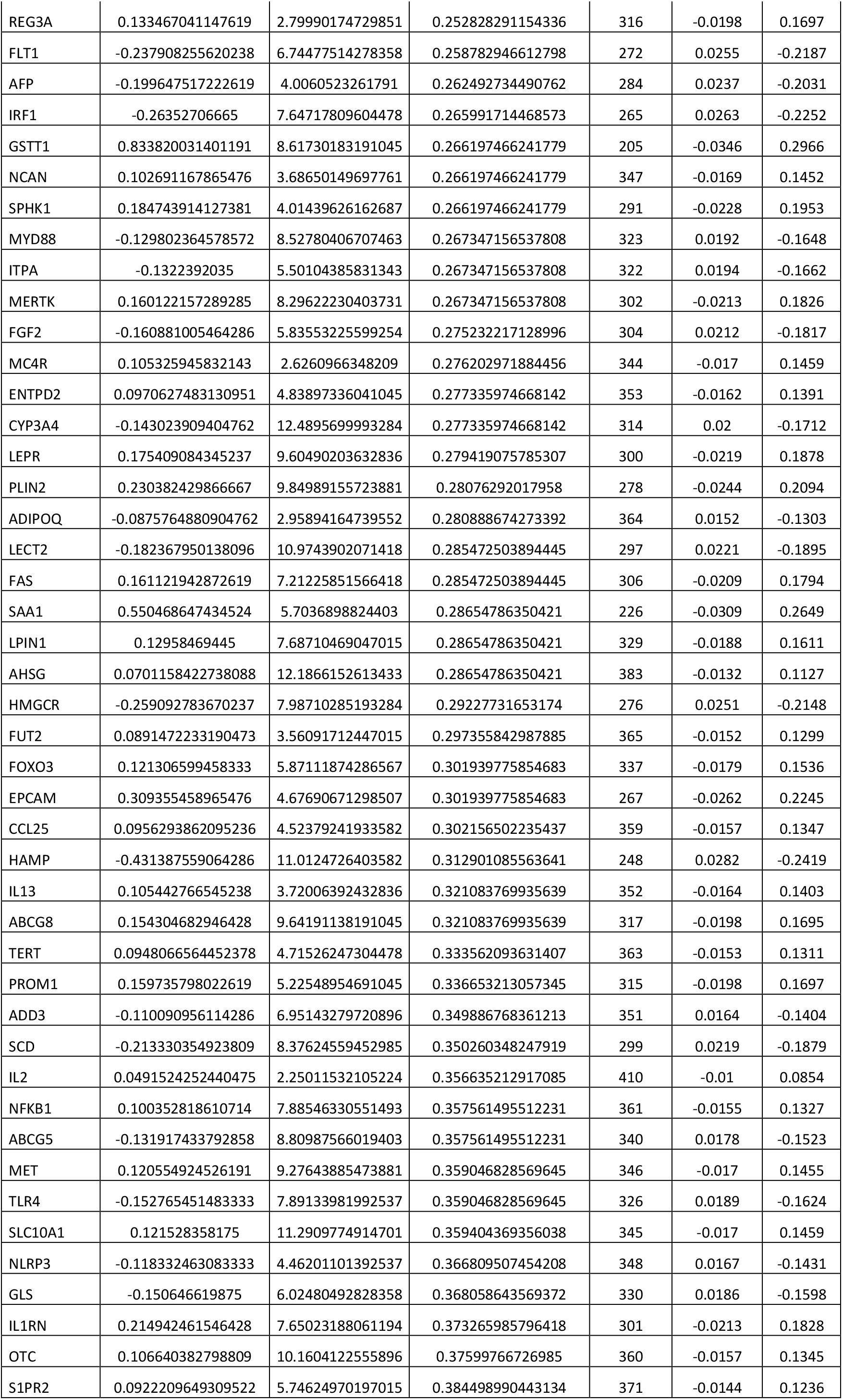
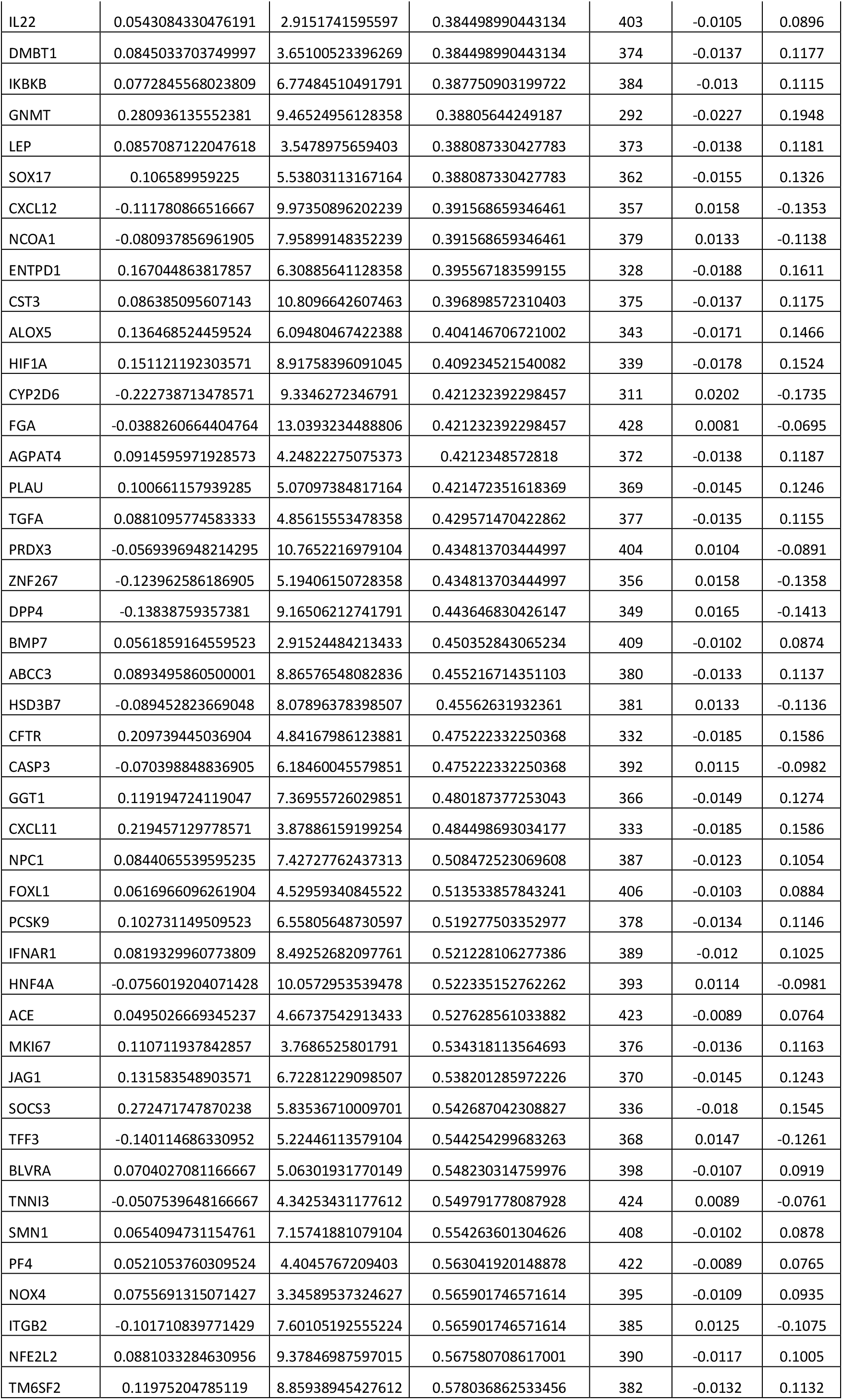
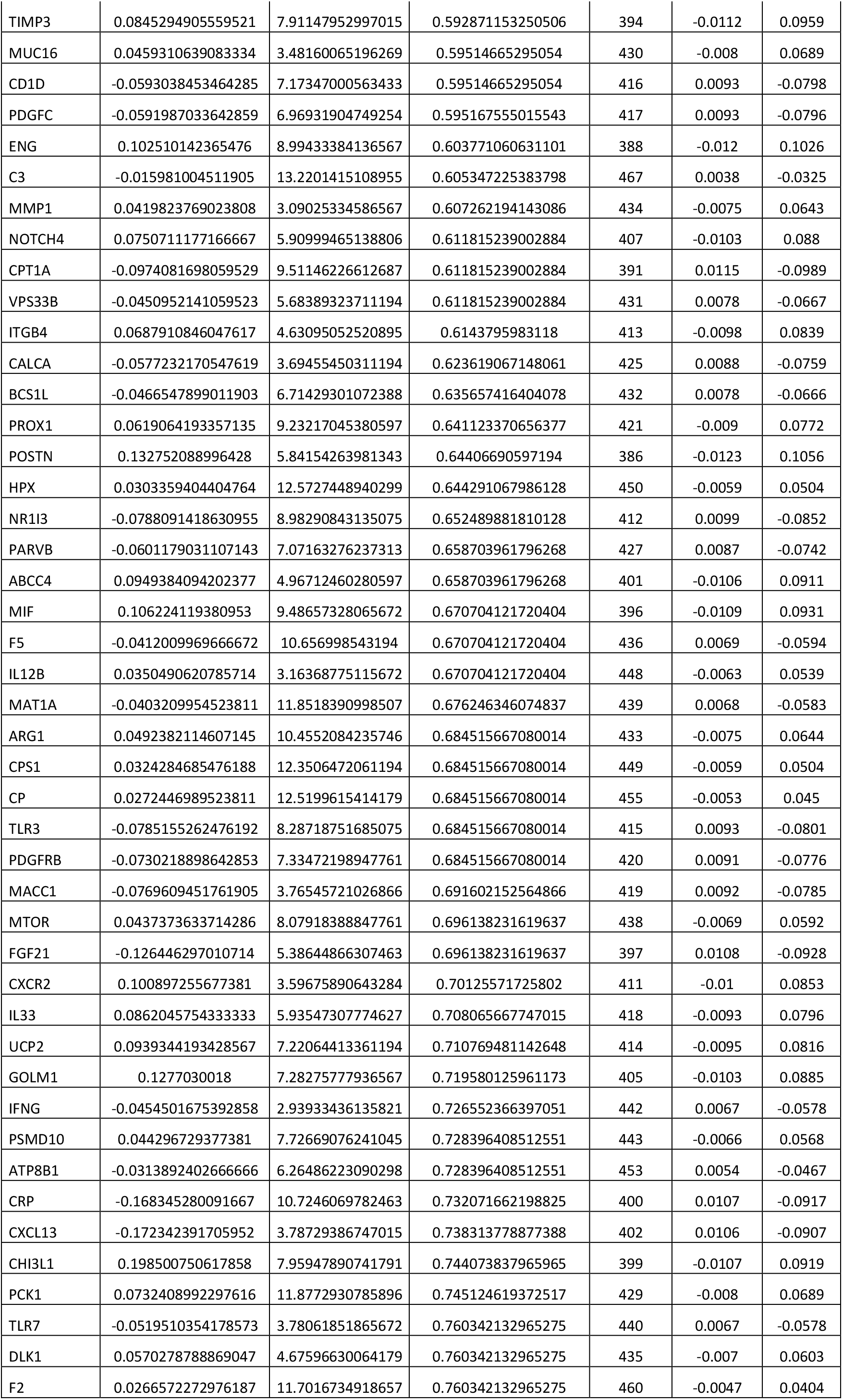
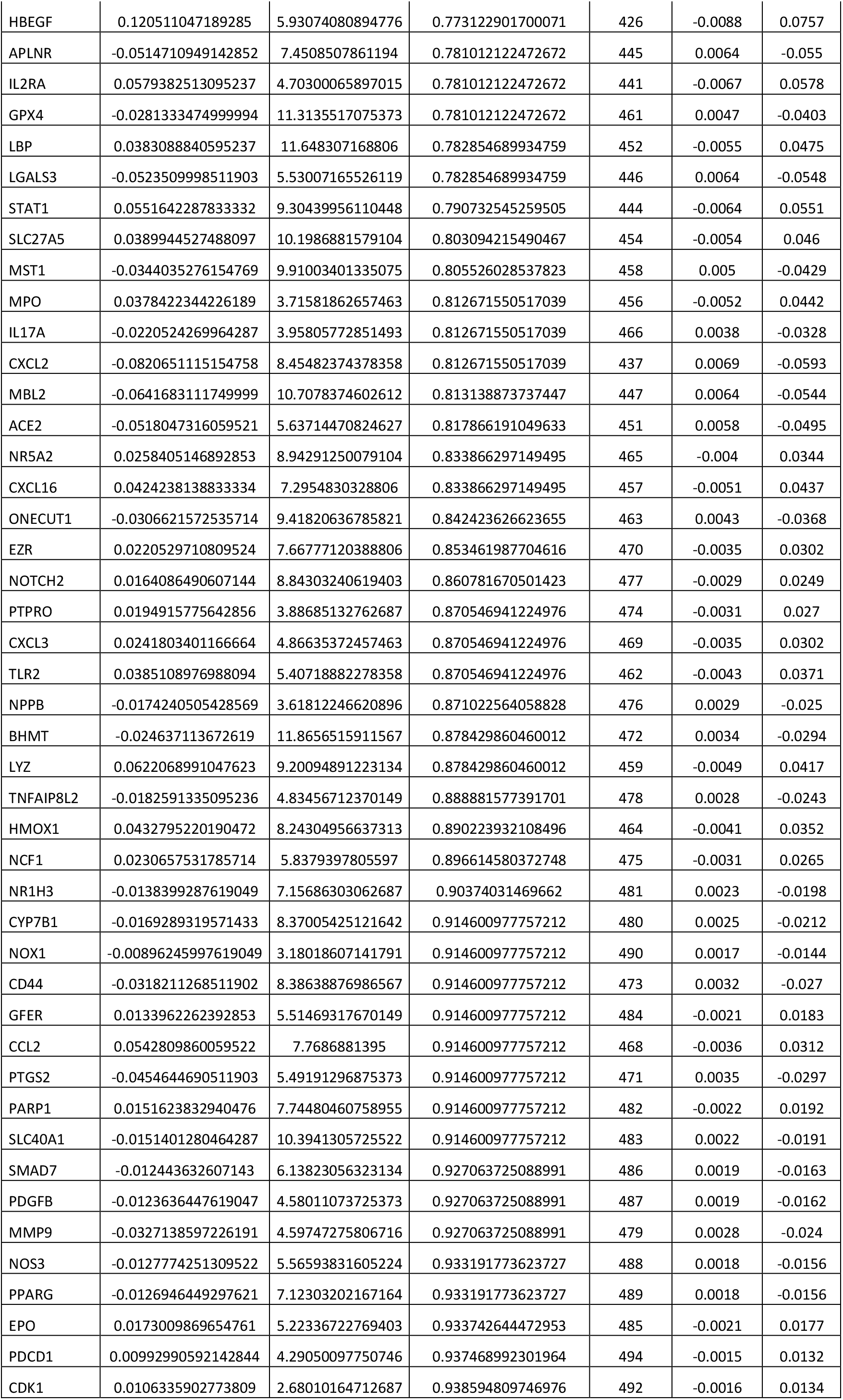
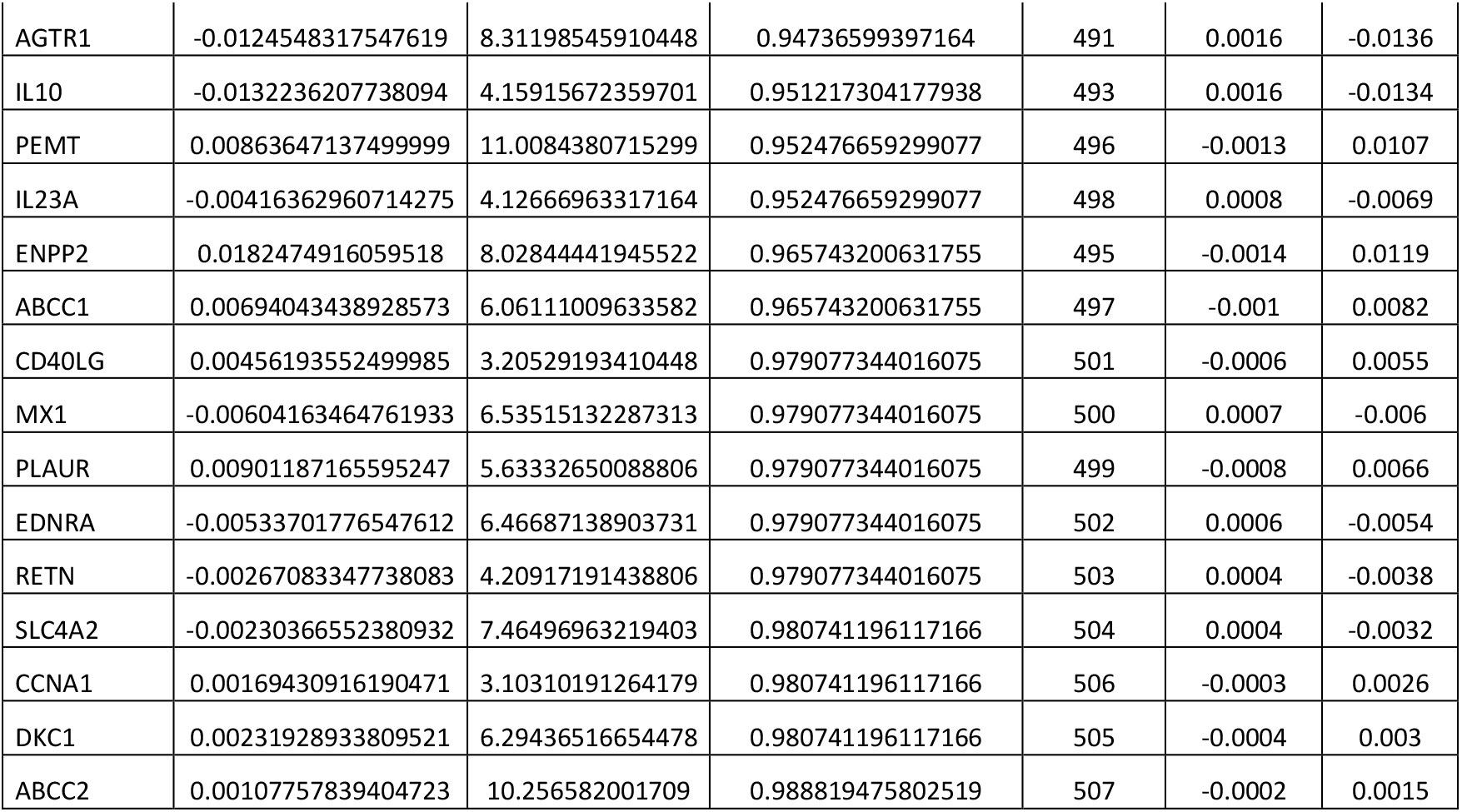
**Symptoms related genes found differentially expressed in liver from primary sclerosing cholangitis patients:** Differential expressed gene analysis results (logarithm Fold Change, Average Epression and Adjust p-values) are presented with respective machine learning predictive score for supervized sample categories of GSE61256 dataset: PSC versus other liver samples)

**Supplemental Table 3:**
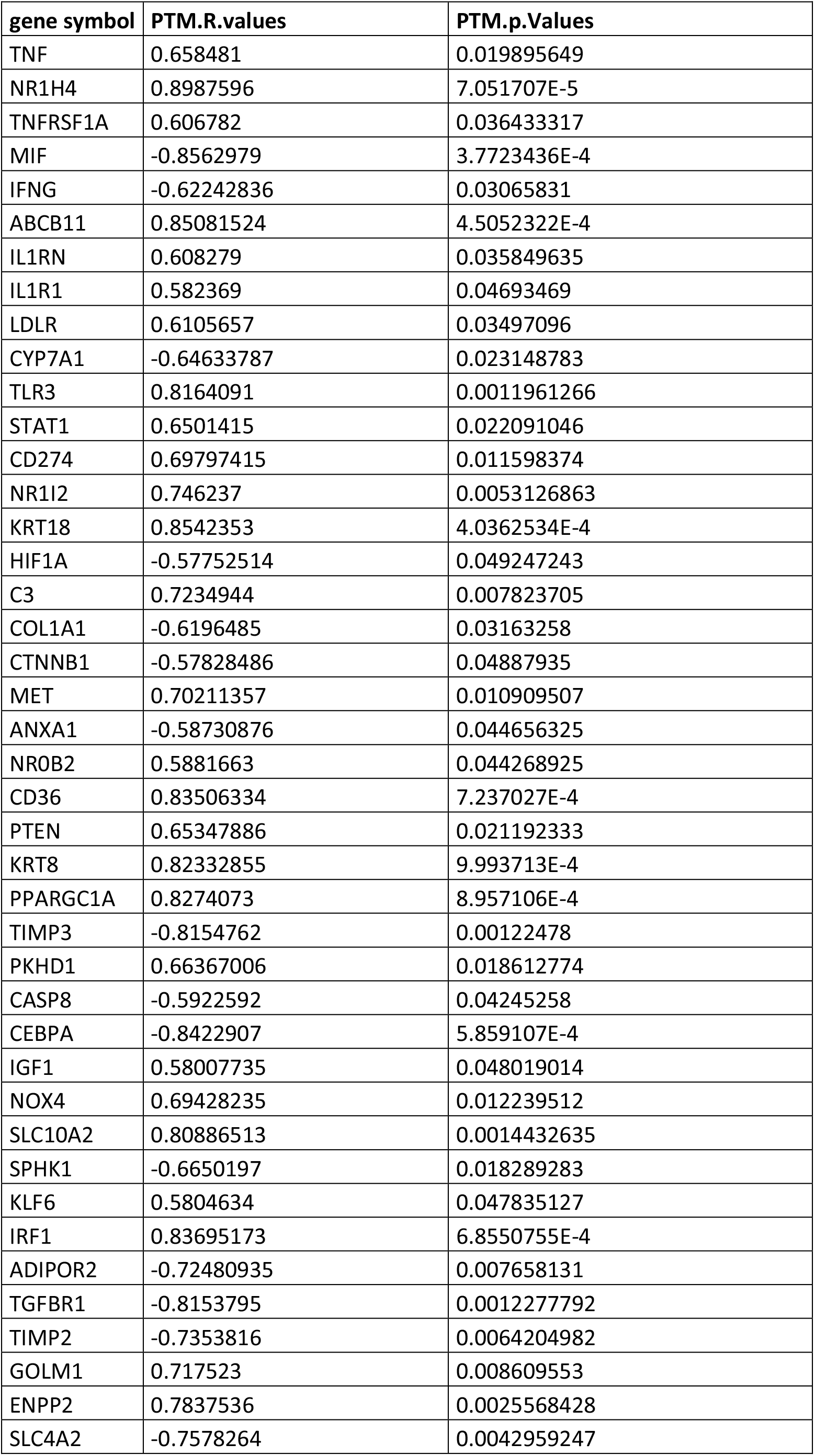
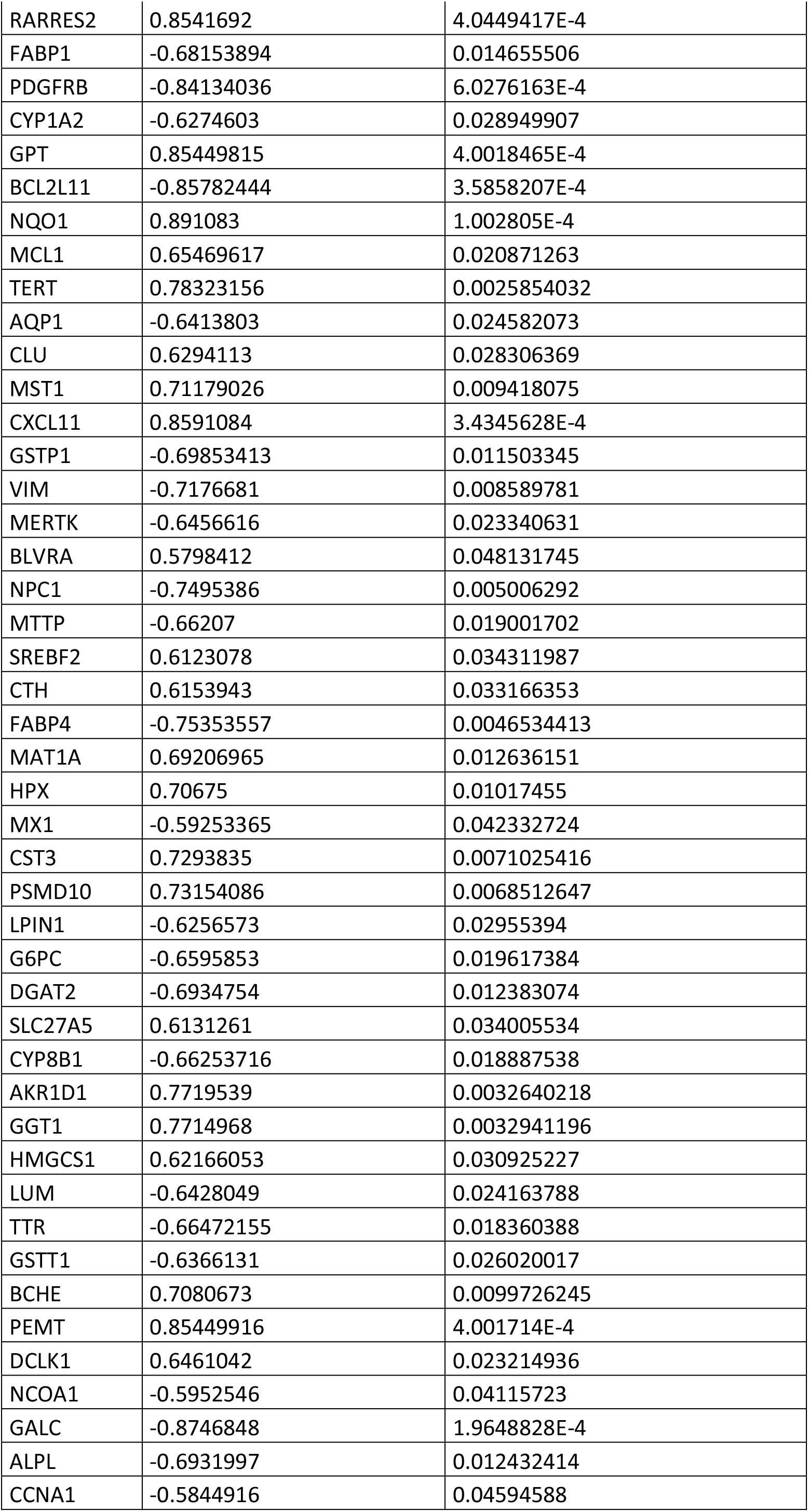
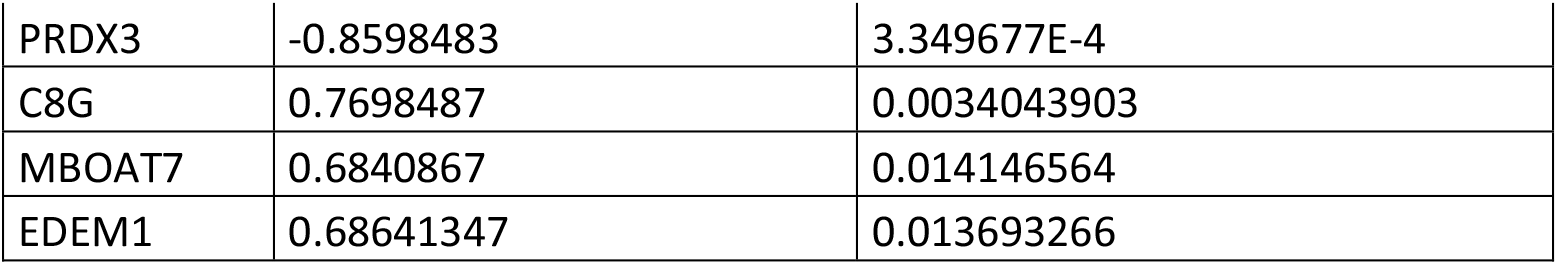
**Pavlidis Template Matching for FXR dependency in liver transcriptome:** Pavlidis template analysis results to describe FXR regulation dependency of PSC genes in liver samples of GSE54557 dataset. For each gene R-Pearson correlation coefficient and its respective p-value are presented in this table.

**Supplemental Table 4:**
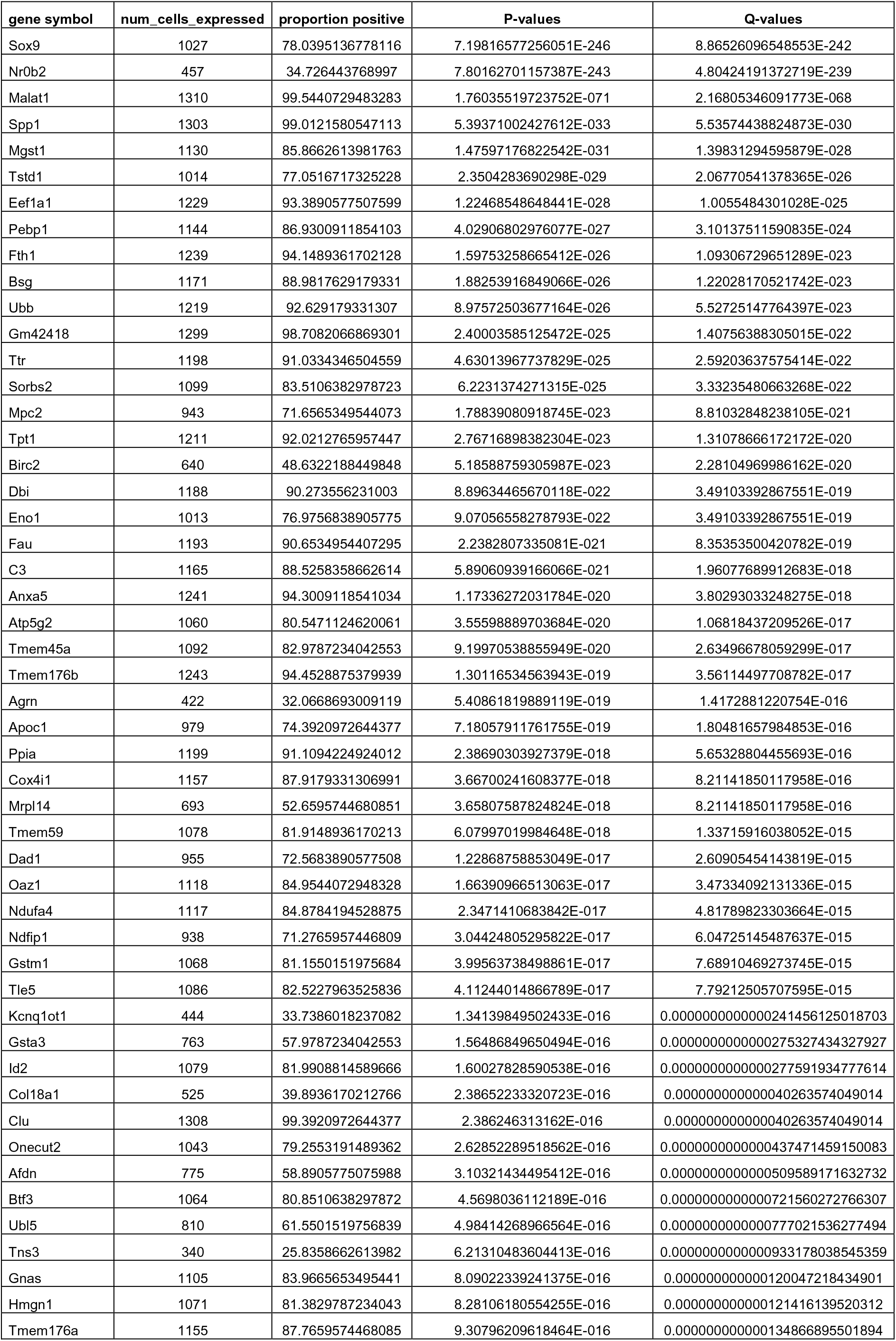
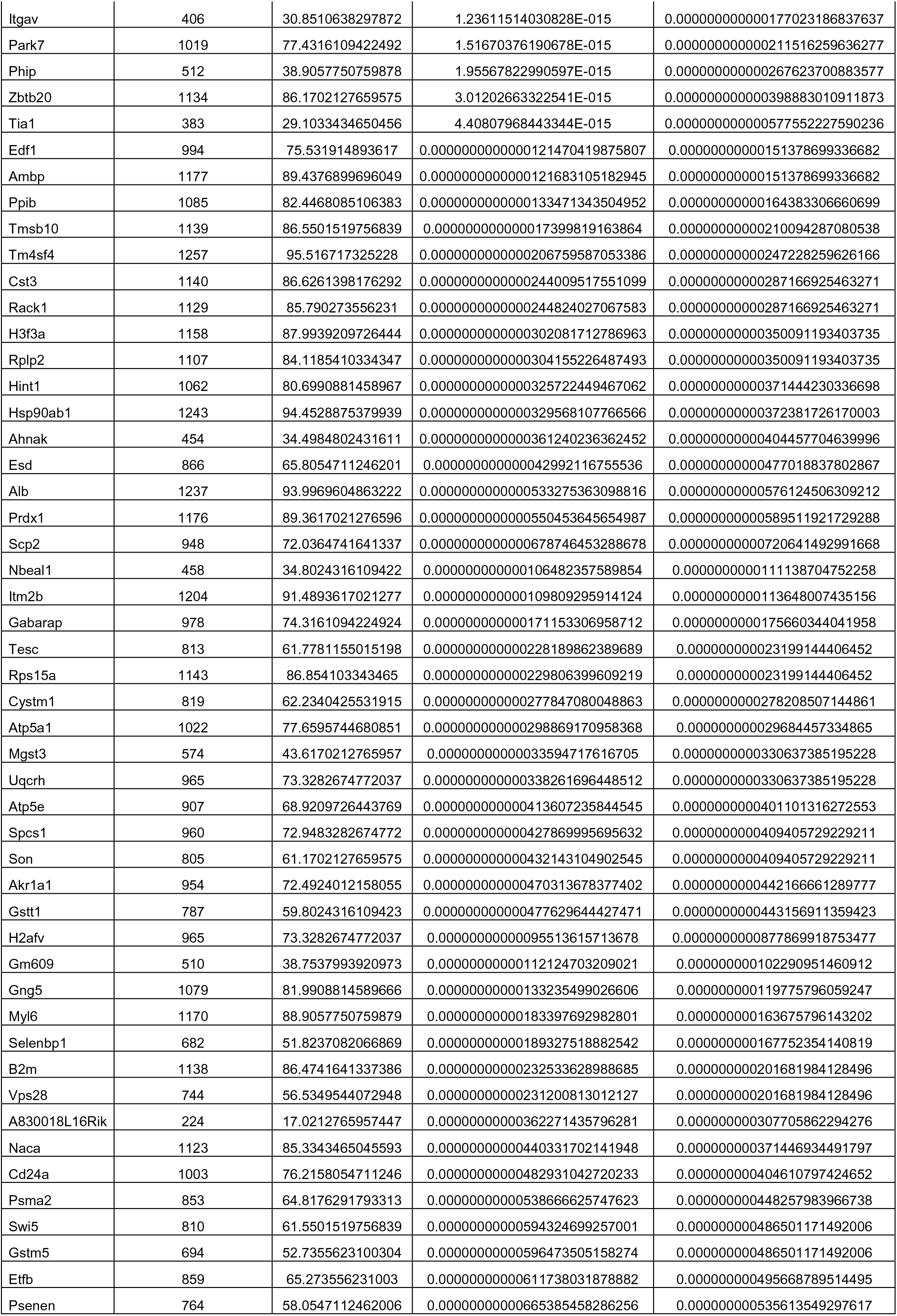
**Best one hundred genes found to be significant pseudotime trajectory of Abcb4-/- cholangiocytes :** Best one hundred ranked genes found as significant on pseudotime cell trajectory of cholangiocytes Abcb4-/- based on alternative expression of Nr0b2 and Sox9 in GSE168758

## REFERENCES

Antoniou, A., Raynaud, P., Cordi, S., Zong, Y., Tronche, F., Stanger, B. Z., … Lemaigre, F. P. (2009). Intrahepatic bile ducts develop according to a new mode of tubulogenesis regulated by the transcription factor SOX9. Gastroenterology, 136(7), 2325-2333. doi: 10.1053/j.gastro.2009.02.051

Benezra, R., Davis, R. L., Lockshon, D., Turner, D. L., & Weintraub, H. (1990). The protein Id : A negative regulator of helix-loop-helix DNA binding proteins. Cell, 61(1), 49-59. doi: 10.1016/0092-8674(90)90214-y

Boonstra, K., van Erpecum, K. J., van Nieuwkerk, K. M. J., Drenth, J. P. H., Poen, A. C., Witteman, B. J. M., … Ponsioen, C. Y. (2012). Primary sclerosing cholangitis is associated with a distinct phenotype of inflammatory bowel disease. Inflammatory Bowel Diseases, 18(12), 2270-2276. doi: 10.1002/ibd.22938

Brandt, C., McFie, P. J., & Stone, S. J. (2016). Biochemical characterization of human acyl coenzyme A : 2-monoacylglycerol acyltransferase-3 (MGAT3). Biochemical and Biophysical Research Communications, 475(3), 264-270. doi: 10.1016/j.bbrc.2016.05.071

Butler, A., Hoffman, P., Smibert, P., Papalexi, E., & Satija, R. (2018). Integrating single-cell transcriptomic data across different conditions, technologies, and species. Nature Biotechnology, 36(5), 411-420. doi: 10.1038/nbt.4096

Chalasani, N., Baluyut, A., Ismail, A., Zaman, A., Sood, G., Ghalib, R., … Hoen, H. (2000). Cholangiocarcinoma in patients with primary sclerosing cholangitis : A multicenter case-control study. Hepatology (Baltimore, Md.), 31(1), 7-11. doi: 10.1002/hep.510310103

Chen, H., Gan, Q., Yang, C., Peng, X., Qin, J., Qiu, S., … Peng, Y. (2019). A novel role of glutathione S-transferase A3 in inhibiting hepatic stellate cell activation and rat hepatic fibrosis. Journal of Translational Medicine, 17(1), 280. doi: 10.1186/s12967-019-2027-8

Chen, J., Bardes, E. E., Aronow, B. J., & Jegga, A. G. (2009). ToppGene Suite for gene list enrichment analysis and candidate gene prioritization. Nucleic Acids Research, 37(Web Server issue), W305–311. doi: 10.1093/nar/gkp427

Cline, M. S., Smoot, M., Cerami, E., Kuchinsky, A., Landys, N., Workman, C., … Bader, G. D. (2007). Integration of biological networks and gene expression data using Cytoscape. Nature Protocols, 2(10), 2366–2382. doi: 10.1038/nprot.2007.324

Crawford, D. R., Ilic, Z., Guest, I., Milne, G. L., Hayes, J. D., & Sell, S. (2017). Characterization of liver injury, oval cell proliferation and cholangiocarcinogenesis in glutathione S-transferase A3 knockout mice. Carcinogenesis, 38(7), 717-727. doi: 10.1093/carcin/bgx048

Desterke, C., Lorenzo, H. K., & Candelier, J.-J. (2021). Text Mining Gene Selection to Understand Pathological Phenotype Using Biological Big Data. In N. Helder I. (Éd.), Bioinformatics. Brisbane (AU): Exon Publications. Consulté à l’adresse http://www.ncbi.nlm.nih.gov/books/NBK569563/

Dwyer, B. J., Jarman, E. J., Gogoi-Tiwari, J., Ferreira-Gonzalez, S., Boulter, L., Guest, R. V., … Forbes, S. J. (2021). TWEAK/Fn14 signalling promotes cholangiocarcinoma niche formation and progression. Journal of Hepatology, 74(4), 860-872. doi: 10.1016/j.jhep.2020.11.018

Fickert, P., Fuchsbichler, A., Wagner, M., Zollner, G., Kaser, A., Tilg, H., … Trauner, M. (2004). Regurgitation of bile acids from leaky bile ducts causes sclerosing cholangitis in Mdr2 (Abcb4) knockout mice. Gastroenterology, 127(1), 261-274. doi: 10.1053/j.gastro.2004.04.009

Fickert, P., Pollheimer, M. J., Beuers, U., Lackner, C., Hirschfield, G., Housset, C., … International PSC Study Group (IPSCSG). (2014). Characterization of animal models for primary sclerosing cholangitis (PSC). Journal of Hepatology, 60(6), 1290–1303. doi: 10.1016/j.jhep.2014.02.006

Flamant, L., Roegiers, E., Pierre, M., Hayez, A., Sterpin, C., De Backer, O., … Michiels, C. (2012). TMEM45A is essential for hypoxia-induced chemoresistance in breast and liver cancer cells. BMC Cancer, 12, 391. doi: 10.1186/1471-2407-12-391

Fleuren, W. W. M., & Alkema, W. (2015). Application of text mining in the biomedical domain. Methods (San Diego, Calif.), 74, 97-106. doi: 10.1016/j.ymeth.2015.01.015

Floreani, A., & De Martin, S. (2021). Treatment of primary sclerosing cholangitis. Digestive and Liver Disease: Official Journal of the Italian Society of Gastroenterology and the Italian Association for the Study of the Liver, S1590-8658(21)00213-9. doi: 10.1016/j.dld.2021.04.028

Fontaine, J.-F., Priller, F., Barbosa-Silva, A., & Andrade-Navarro, M. A. (2011). Génie : Literature-based gene prioritization at multi genomic scale. Nucleic Acids Research, 39(Web Server issue), W455-461. doi: 10.1093/nar/gkr246

The Gene Ontology Consortium. (2017). Expansion of the Gene Ontology knowledgebase and resources. Nucleic Acids Research, 45(D1), D331–D338. doi: 10.1093/nar/gkw1108

Goodwin, B., Jones, S. A., Price, R. R., Watson, M. A., McKee, D. D., Moore, L. B., … Kliewer, S. A. (2000). A regulatory cascade of the nuclear receptors FXR, SHP-1, and LRH-1 represses bile acid biosynthesis. Molecular Cell, 6(3), 517–526. doi: 10.1016/s1097-2765(00)00051-4

Gronemeyer, H., Gustafsson, J.-A., & Laudet, V. (2004). Principles for modulation of the nuclear receptor superfamily. Nature Reviews. Drug Discovery, 3(11), 950-964. doi: 10.1038/nrd1551

Guo, J., Chen, L., Luo, N., Yang, W., Qu, X., & Cheng, Z. (2015). Inhibition of TMEM45A suppresses proliferation, induces cell cycle arrest and reduces cell invasion in human ovarian cancer cells. Oncology Reports, 33(6), 3124-3130. doi: 10.3892/or.2015.3902

Hang, S., Paik, D., Yao, L., Kim, E., Trinath, J., Lu, J., … Huh, J. R. (2019). Bile acid metabolites control TH17 and Treg cell differentiation. Nature, 576(7785), 143-148. doi: 10.1038/s41586-019-1785-z

Harder, J., Müller, M. J., Fuchs, M., Gumpp, V., Schmitt-Graeff, A., Fischer, R., … Hasskarl, J. (2013). Inhibitor of differentiation proteins do not influence prognosis of biliary tract cancer. World Journal of Gastroenterology, 19(48), 9334-9342. doi: 10.3748/wjg.v19.i48.9334

Horvath, S., Erhart, W., Brosch, M., Ammerpohl, O., von Schönfels, W., Ahrens, M., … Hampe, J. (2014). Obesity accelerates epigenetic aging of human liver. Proceedings of the National Academy of Sciences of the United States of America, 111(43), 15538-15543. doi: 10.1073/pnas.1412759111

Inagaki, T., Choi, M., Moschetta, A., Peng, L., Cummins, C. L., McDonald, J. G., … Kliewer, S. A. (2005). Fibroblast growth factor 15 functions as an enterohepatic signal to regulate bile acid homeostasis. Cell Metabolism, 2(4), 217-225. doi: 10.1016/j.cmet.2005.09.001

Johansson, A. S., & Mannervik, B. (2001). Human glutathione transferase A3-3, a highly efficient catalyst of double-bond isomerization in the biosynthetic pathway of steroid hormones. The Journal of Biological Chemistry, 276(35), 33061-33065. doi: 10.1074/jbc.M104539200

Karlsen, T. H., Folseraas, T., Thorburn, D., & Vesterhus, M. (2017). Primary sclerosing cholangitis—A comprehensive review. Journal of Hepatology, 67(6), 1298-1323. doi: 10.1016/j.jhep.2017.07.022

Krallinger, M., Valencia, A., & Hirschman, L. (2008). Linking genes to literature : Text mining, information extraction, and retrieval applications for biology. Genome Biology, 9 *Suppl 2*, S8. doi: 10.1186/gb-2008-9-s2-s8

Lazaridis, K. N., & LaRusso, N. F. (2016). Primary Sclerosing Cholangitis. The New England Journal of Medicine, 375(12), 1161-1170. doi: 10.1056/NEJMra1506330

Lê, S., Josse, J., & Husson, F. (2008). FactoMineR: An R Package for Multivariate Analysis. Journal of Statistical Software, 25(1), 1-18.

Lee, S., Stewart, S., Nagtegaal, I., Luo, J., Wu, Y., Colditz, G., … Allred, D. C. (2012). Differentially expressed genes regulating the progression of ductal carcinoma in situ to invasive breast cancer. Cancer Research, 72(17), 4574-4586. doi: 10.1158/0008-5472.CAN-12-0636

Manawapat-Klopfer, A., Thomsen, L. T., Martus, P., Munk, C., Russ, R., Gmuender, H., … Iftner, T. (2016). TMEM45A, SERPINB5 and p16INK4A transcript levels are predictive for development of high-grade cervical lesions. American Journal of Cancer Research, 6(7), 1524–1536.

Mertz, A., Nguyen, N. A., Katsanos, K. H., & Kwok, R. M. (2019). Primary sclerosing cholangitis and inflammatory bowel disease comorbidity : An update of the evidence. Annals of Gastroenterology, 32(2), 124-133. doi: 10.20524/aog.2019.0344

Molodecky, N. A., Kareemi, H., Parab, R., Barkema, H. W., Quan, H., Myers, R. P., & Kaplan, G. G. (2011). Incidence of primary sclerosing cholangitis : A systematic review and meta-analysis. Hepatology (Baltimore, Md.), 53(5), 1590-1599. doi: 10.1002/hep.24247

Ozdil, B., Cosar, A., Akkiz, H., Sandikci, M., & Kece, C. (2011). New therapeutic option with N-acetylcysteine for primary sclerosing cholangitis : Two case reports. American Journal of Therapeutics, 18(3), e71–74. doi: 10.1097/MJT.0b013e3181c42758

Pavlidis, P., & Noble, W. S. (2001). Analysis of strain and regional variation in gene expression in mouse brain. Genome Biology, 2(10), RESEARCH0042.

Pirola, C. J., & Sookoian, S. (2021). The lipidome in nonalcoholic fatty liver disease : Actionable targets. Journal of Lipid Research, 62, 100073. doi: 10.1016/j.jlr.2021.100073

Reich, M., Spomer, L., Klindt, C., Fuchs, K., Stindt, J., Deutschmann, K., … Keitel, V. (2021). Downregulation of TGR5 (GPBAR1) in biliary epithelial cells contributes to the pathogenesis of sclerosing cholangitis. *Journal of Hepatology*, S0168–8278(21)00244-0. doi: 10.1016/j.jhep.2021.03.029

Ritchie, M. E., Phipson, B., Wu, D., Hu, Y., Law, C. W., Shi, W., & Smyth, G. K. (2015). Limma powers differential expression analyses for RNA-sequencing and microarray studies. Nucleic Acids Research, 43(7), e47. doi: 10.1093/nar/gkv007

Rizvi, S., Eaton, J. E., & Gores, G. J. (2015). Primary Sclerosing Cholangitis as a Premalignant Biliary Tract Disease : Surveillance and Management. Clinical Gastroenterology and Hepatology: The Official Clinical Practice Journal of the American Gastroenterological Association, 13(12), 2152-2165. doi: 10.1016/j.cgh.2015.05.035

Rühlemann, M., Liwinski, T., Heinsen, F.-A., Bang, C., Zenouzi, R., Kummen, M., … Franke, A. (2019). Consistent alterations in faecal microbiomes of patients with primary sclerosing cholangitis independent of associated colitis. Alimentary Pharmacology & Therapeutics, 50(5), 580-589. doi: 10.1111/apt.15375

Sabino, J., Vieira-Silva, S., Machiels, K., Joossens, M., Falony, G., Ballet, V., … Raes, J. (2016). Primary sclerosing cholangitis is characterised by intestinal dysbiosis independent from IBD. Gut, 65(10), 1681-1689. doi: 10.1136/gutjnl-2015-311004

Sakaguchi, S., Miyara, M., Costantino, C. M., & Hafler, D. A. (2010). FOXP3+ regulatory T cells in the human immune system. Nature Reviews. Immunology, 10(7), 490-500. doi: 10.1038/nri2785

Sayers, E. (2010). Entrez Programming Utilities Help (https://www.ncbi.nlm.nih.gov/books/NBK25501/). National Center for Biotechnology Information (US). (https://www.ncbi.nlm.nih.gov/books/NBK25501/). Consulté à l’adresse https://www.ncbi.nlm.nih.gov/books/NBK25501/

Sayers, E. W., Barrett, T., Benson, D. A., Bryant, S. H., Canese, K., Chetvernin, V., … Ye, J. (2009). Database resources of the National Center for Biotechnology Information. Nucleic Acids Research, 37(Database issue), D5-15. doi: 10.1093/nar/gkn741

Sebode, M., Peiseler, M., Franke, B., Schwinge, D., Schoknecht, T., Wortmann, F., … Schramm, C. (2014). Reduced FOXP3(+) regulatory T cells in patients with primary sclerosing cholangitis are associated with IL2RA gene polymorphisms. Journal of Hepatology, 60(5), 1010-1016. doi: 10.1016/j.jhep.2013.12.027

Seol, W., Choi, H. S., & Moore, D. D. (1996). An orphan nuclear hormone receptor that lacks a DNA binding domain and heterodimerizes with other receptors. Science (New York, N.Y.), 272(5266), 1336-1339. doi: 10.1126/science.272.5266.1336

Sinal, C. J., Tohkin, M., Miyata, M., Ward, J. M., Lambert, G., & Gonzalez, F. J. (2000). Targeted disruption of the nuclear receptor FXR/BAR impairs bile acid and lipid homeostasis. Cell, 102(6), 731-744. doi: 10.1016/s0092-8674(00)00062-3

Subramanian, A., Tamayo, P., Mootha, V. K., Mukherjee, S., Ebert, B. L., Gillette, M. A., … Mesirov, J. P. (2005). Gene set enrichment analysis : A knowledge-based approach for interpreting genome-wide expression profiles. Proceedings of the National Academy of Sciences of the United States of America, 102(43), 15545-15550. doi: 10.1073/pnas.0506580102

Sun, W., Qiu, G., Zou, Y., Cai, Z., Wang, P., Lin, X., … Hu, G. (2015). Knockdown of TMEM45A inhibits the proliferation, migration and invasion of glioma cells. International Journal of Clinical and Experimental Pathology, 8(10), 12657-12667.

Szklarczyk, D., Morris, J. H., Cook, H., Kuhn, M., Wyder, S., Simonovic, M., … von Mering, C. (2017). The STRING database in 2017 : Quality-controlled protein-protein association networks, made broadly accessible. Nucleic Acids Research, 45(D1), D362-D368. doi: 10.1093/nar/gkw937

Tahboub Amawi, A. D., Tremaine, W. J., & Venkatesh, S. K. (2020). Pembrolizumab-Induced Sclerosing Cholangitis. Clinical Gastroenterology and Hepatology: The Official Clinical Practice Journal of the American Gastroenterological Association, S1542-3565(20)31638-4. doi: 10.1016/j.cgh.2020.11.048

Tietz-Bogert, P. S., Kim, M., Cheung, A., Tabibian, J. H., Heimbach, J. K., Rosen, C. B., … O’Hara, S. P. (2018). Metabolomic Profiling of Portal Blood and Bile Reveals Metabolic Signatures of Primary Sclerosing Cholangitis. International Journal of Molecular Sciences, 19(10), E3188. doi: 10.3390/ijms19103188

Trapnell, C., Cacchiarelli, D., Grimsby, J., Pokharel, P., Li, S., Morse, M., … Rinn, J. L. (2014). The dynamics and regulators of cell fate decisions are revealed by pseudotemporal ordering of single cells. Nature Biotechnology, 32(4), 381-386. doi: 10.1038/nbt.2859

Vestentoft, P. S., Jelnes, P., Hopkinson, B. M., Vainer, B., Møllgård, K., Quistorff, B., & Bisgaard, H. C. (2011). Three-dimensional reconstructions of intrahepatic bile duct tubulogenesis in human liver. BMC Developmental Biology, 11, 56. doi: 10.1186/1471-213X-11-56

Wickham, H. (2009). ggplot2 : Elegant Graphics for Data Analysis. Springer-Verlag *New York*.

Wrzesiński, T., Szelag, M., Cieślikowski, W. A., Ida, A., Giles, R., Zodro, E., … Wesoly, J. (2015). Expression of pre-selected TMEMs with predicted ER localization as potential classifiers of ccRCC tumors. BMC Cancer, 15, 518. doi: 10.1186/s12885-015-1530-4

Xia, J., Gill, E. E., & Hancock, R. E. W. (2015). NetworkAnalyst for statistical, visual and network-based meta-analysis of gene expression data. Nature Protocols, 10(6), 823-844. doi: 10.1038/nprot.2015.052

Xu, C., & Su, Z. (2015). Identification of cell types from single-cell transcriptomes using a novel clustering method. Bioinformatics (Oxford, England), 31(12), 1974-1980. doi: 10.1093/bioinformatics/btv088

Yang, Y., Sharma, R., Zimniak, P., & Awasthi, Y. C. (2002). Role of alpha class glutathione S-transferases as antioxidant enzymes in rodent tissues. Toxicology and Applied Pharmacology, 182(2), 105-115. doi: 10.1006/taap.2002.9450

Zhan, L., Liu, H.-X., Fang, Y., Kong, B., He, Y., Zhong, X.-B., … Guo, G. L. (2014). Genome-wide binding and transcriptome analysis of human farnesoid X receptor in primary human hepatocytes. PloS One, 9(9), e105930. doi: 10.1371/journal.pone.0105930

Zhang, Y., Hagedorn, C. H., & Wang, L. (2011). Role of nuclear receptor SHP in metabolism and cancer. Biochimica Et Biophysica Acta, 1812(8), 893-908. doi: 10.1016/j.bbadis.2010.10.006

